# Multifaceted adaptation of the neural decision process with prior knowledge of time constraints and stimulus probability

**DOI:** 10.1101/715318

**Authors:** Simon P. Kelly, Elaine A. Corbett, Redmond G. O’Connell

## Abstract

When selecting actions in response to noisy sensory stimuli, the brain can exploit prior knowledge of time constraints, stimulus discriminability and stimulus probability to hone the decision process. Although behavioral models typically explain such effects through adjustments to decision criteria only, the full range of underlying neural process adjustments remains to be established. Here, we draw on human neurophysiological signals reflecting decision formation to construct and constrain a multi-tiered model of prior-informed motion discrimination, in which a motor-independent representation of cumulative evidence feeds build-to-threshold motor signals that receive additional dynamic urgency and bias signal components. The neurally-informed model not only provides a superior quantitative fit to prior-biased behavior across three distinct task regimes (easy, time-pressured and weak evidence), but also reveals adjustments to evidence accumulation rate, urgency rate, and the timing of accumulation onset and motor execution which go undetected or are discrepant in more standard diffusion-model analysis of behavior.

## Introduction

Our ability to adapt our decision processes to take account of time constraints and relative stimulus probability is critical to behaving effectively in diverse environments. When time is short, we can expedite our decisions but usually at a cost to accuracy, and when informative cues are available, we tend to bias our decisions in favour of the choice alternative we know is more likely to be correct. Computational decision models based on the principle of bounded evidence accumulation (Smith and Ratcliff 2004; Link and Heath 1975) offer elegant explanations of each of these behavioral phenomena through single parameter adjustments, which have been borne out in a vast number of behavioral modeling studies. Specifically, it is almost unanimously found that accuracy is traded for speed by lowering the quantity of evidence required to trigger commitment to either alternative (i.e., narrowing the decision bounds), while behavioral biases are effected by starting the accumulation process closer to the bound for the more probable alternative (Ratcliff and McKoon 2008; Heitz 2014; Mulder et al. 2012). There now also exists substantial neurophysiological evidence for these bound and starting point adjustments (Bogacz et al. 2010; Ivanoff et al. 2008; Forstmann et al. 2008; Hanks et al. 2014; Rorie et al. 2010; de Lange et al. 2013; van Veen et al. 2008). However, while these dominant adjustments are usually sufficient to capture the aggregate behavioral phenomena observed, it is becoming increasingly apparent that they are unlikely to be the only adjustments made (Heitz and Schall 2013; Hanks et al. 2011; Hanks et al. 2014; Rae et al. 2014).

Some behavioral modeling studies have suggested that, in addition to lowered decision bounds, speed pressure induces a shortening of “non-decision time,” a parameter used to capture the additional delays associated with stimulus encoding and motor execution that are not attributable to the deliberative decision process itself (Voss et al. 2004; Rinkenauer et al. 2004; Rae et al. 2014; Arnold et al. 2015). Other work has suggested that speed pressure leads to a reduction in the mean rate of accumulation, captured in “drift rate” (formally, the statistical expectation of the evidence; (Rae et al. 2014; Dutilh et al. 2018). Meanwhile, effects of prior stimulus probability have been observed on drift rate in addition to starting point (Dunovan et al. 2014). However, adjustments to these parameters have only been observed in a small minority of studies and, due to a lack of strong constraints, their presence or absence can be extremely difficult to reliably establish based on behavioral data alone (Dutilh et al. 2018). Yet, establishing the full range of parameter adjustments is critical to fully understanding the capacity and mechanisms for adapting our decisions to the environment, and how they may be disrupted by clinical brain disorders.

Alongside this increasing scrutiny of decision process adaptations, cognitive modeling and neurophysiological research has recently trained a spotlight on the validity of certain core assumptions of standard sequential sampling models. In particular, there has been an escalating debate regarding whether subjects adopt bounds that are constant for the duration of each decision as prescribed in standard models (Ratcliff et al. 2016), or alternatively, decrease them over time within a trial (Hanks et al. 2014; Thura et al. 2012; Hawkins et al. 2015; Trueblood et al. 2019). Such a collapsing-bound scheme has been established in theoretical work to be optimal when sensory discriminability varies across trials (Malhotra et al. 2018; Moran 2015), when there is a cost to continued accumulation (Drugowitsch et al. 2012; Boehm et al. 2019), or when missed deadlines are penalised (Frazier and Yu 2008). Yet, behavioral modeling evidence is extremely mixed as to whether human subjects actually collapse their decision bounds in practice (Malhotra et al. 2017; Evans and Hawkins 2019; Palestro et al. 2018), with several recent formal model comparisons clearly favouring standard constant-bound models (Hawkins et al. 2015; Voskuilen et al. 2016; Evans, Hawkins, et al. 2019). In fact, definitively establishing the role of collapsing bounds based solely on behavioral modeling is highly challenging because one of its primary qualitative expressions – increased error rates on trials with longer RTs – can be alternatively produced with increased between-trial drift rate variability (Ratcliff et al. 2016), and many collapsing bound models suffer from identifiability issues due to a lack of constraints (Evans, Trueblood, et al. 2019). However, there is increasing recognition that such impasses can be broken by also considering the unique predictions that competing decision models make regarding the neural processes underlying decision formation (Turner et al. 2016; Purcell and Palmeri 2017; O’Connell et al. 2018).

Neurophysiology studies of the monkey, rat and human brain have established that the evidence-dependent build-to-threshold dynamics predicted by accumulation-to-bound models are reflected in neural signals in several brain circuits, especially those associated with the planning of decision-reporting actions (Hanes and Schall 1996; Roitman and Shadlen 2002; Gold and Shadlen 2007; Donner et al. 2009; van Vugt et al. 2014; O’Connell et al. 2012; Hanks and Summerfield 2017). In fact, several such studies have uncovered evidence for a temporally increasing, evidence-independent ‘urgency’ component of the build-up in these signals that provides a neural mechanism for implementing collapsing bounds (Churchland et al. 2008; Hanks et al. 2014; Murphy et al. 2016; Steinemann et al. 2018). In particular, when one study implemented a model with an urgency function that was constrained to match neural urgency measurements, it produced excellent fits to behavior (Hanks et al. 2014). Thus, there is strong but as yet underexploited potential for neurophysiological decision signals that are finely resolved in time to guide the construction as well as constrain the fitting of models whose structure adheres to the underlying neural implementation at a systems level (O’Connell et al. 2018).

In the present study we develop and implement a multi-level accumulation-to-bound model that can simultaneously explain the choice-predictive dynamics exhibited by neural signatures of decision formation as well as behavior across a range of task conditions manipulating the prior knowledge of time constraints, stimulus discriminability and stimulus probability. To do this, we leveraged two functionally distinct decision signals that have been isolated and characterised in human electrophysiology (EEG): premotor Mu/Beta-band activity which reflects the translation of the emerging decision into a specific motor plan (left vs right hand button push; Donner et al. 2009; de Lange et al. 2013; O’Connell et al. 2012; Murphy et al. 2016; Steinemann et al. 2018) and a supramodal, motor-independent signal (centro-parietal positivity, CPP) that builds with cumulative evidence irrespective of the sensory feature or modality or the motor requirements for reporting the decisions (O’Connell et al. 2012; Kelly and O’Connell 2013; Twomey et al. 2016). These two dynamic neural decision signals together provided a rich source of qualitative effects to guide the neurally-informed model’s construction, quantitative neural measurements to constrain key parameter values, and additional dynamic signal features through which key model predictions could be further tested. To briefly summarize, based on distinct modulation patterns observed at the outset and the end of the decision process in the two neural signals, we constructed a model in which sensory evidence accumulation reflected in the CPP feeds downstream motor preparation signals reflected in Mu/Beta-band activity, which also receives additive urgency signal components modulated by temporal/perceptual demands and priors. Meanwhile, measurements of motor preparation magnitudes and latencies were used to directly constrain starting points with respect to fixed motor thresholds and the duration of “non-decision” motor execution processes.

When compared to a standard drift diffusion model (DDM), this neurally-informed (NI) model was found to provide a superior fit to the behavioral data and revealed several novel adjustments made by the subjects based on their prior knowledge, each of which was mirrored in model-independent neural observations. 1) Whereas the DDM suggested that drift rate was reduced under speed pressure for a given evidence strength, the NI model fits to behavior pointed to significantly increased drift rate, which was supported by increased CPP buildup rate. 2) Whereas the DDM specifies a unitary non-decision time parameter, the neural data and NI model enabled the decomposition of this parameter into three distinct components (accumulation onset, evidence encoding onset, post-commitment motor execution), and reveal that they can undergo independent strategic adaptations. 3) Whereas the DDM assumed there was no dynamic urgency component (i.e., no bound collapse), the NI model revealed that dynamic urgency was applied in all regimes in addition to starting point shifts, and that paradoxically, the rate of urgency buildup was steepest under the least urgent regime and shallowest in the most urgent regime – a pattern also reflected in motor preparation signals. 4) Whereas model selection based on the DDM failed to find evidence for modulations of drift rate with prior probability, the NI model indicated the presence of such modulations less ambiguously, and this was further supported by qualitative signatures of drift bias in both behavioral accuracy and in CPP amplitude modulation patterns.

These findings, taken together, establish that decision makers make multiple, sometimes non-intuitive, adjustments to their decision processes to account for prior knowledge of task demands and probabilities. They also affirm the potential explanatory gains offered by using neural signatures of decision formation to not only complement or provide neural correlates for parameters of existing and/or standard models, but also to inform the construction of new, more complex models that capture, at a systems level, the multitiered neural architecture that implements these decision mechanisms.

## Results

Scalp electroencephalographic (EEG) signals were recorded while twenty human participants performed a prior-cued motion discrimination task under three different blocked task regimes: In the ‘Easy’ regime, motion strength was high (20% coherence) and subjects were given a response deadline of 1600ms i.e. the full coherent-motion duration; In the ‘LoCoh’ regime, the same deadline of 1600 ms was again imposed but coherence was close to individual discrimination thresholds (mean ± s.d. 8.4 ± 1.7 %; titrated for each individual); In the ‘Deadline’ regime, coherence was set at the easy level of 20% but with a shorter deadline that was titrated for each individual (mean 485 ± 61 ms). In each trial, subjects first viewed incoherently moving grey dots, which changed after 647 ms to one of three possible colors to indicate that leftward motion was three times more likely, equally likely, or three times less likely than rightward motion, respectively (Figure 1A). After a further 764 ms the dots began to move coherently, always with the same fixed motion strength for a given block. Subjects were asked to respond with their left hand for leftward motion or with their right hand for rightward motion in order to earn points for correct decisions reported within the deadline (0 points received for missed deadlines). The fixed timing meant that subjects could predict evidence onset quite well even under weak evidence. The seamless transition from zero to non-zero coherence at evidence onset avoids non-task specific evoked potentials caused by sudden luminance increments, which would otherwise overlap with and obscure the relevant decision signals (See also Loughnane et al. 2016; Kelly and O’Connell 2013; Twomey et al. 2016).

**Figure 1:**
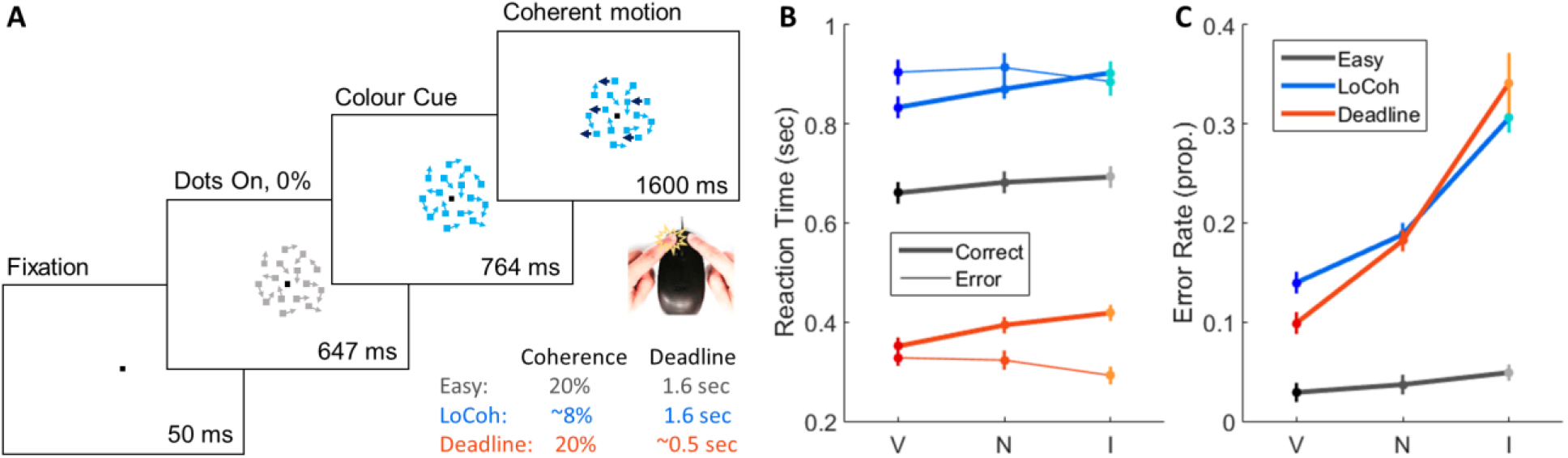
Prior-cued motion discrimination task and behavioral data. A) Trial structure and mode of responding by mouse-button click. B) Mean response time (RT) for valid (V), neutral (N) and invalid (I) trials of each Regime. C) Mean error rate as a function of Regime and Validity. Error bars indicate S.E.M. after between-subject variance is factored out.

Behavior was strongly influenced by both the prevailing perceptual/urgency regimes and the prior information provided by the color-change cues (Figure 1B,C). Response times (RT) were generally much faster in the Deadline regime and much slower in the LoCoh regime, relative to Easy (main effect of Regime: F(2, 38) = 218, p < 1e-323, all pairwise comparisons p<1e-6), and error rate was strongly increased not only due to reduced coherence (main effect of Regime: F(2, 38) = 95.5, p = 1.55e-15; LoCoh > Easy: t(19) = 16.6, p = 4.4e-13) but also due to imposed time pressure (Deadline > Easy: t(19) = 10.6, p = 9.7e-10). Prior cues biased both RT and accuracy in all Regimes, with greater accuracy and faster correct responses in validly-cued trials relative to invalid trials, and neutral trials lying in between (Fig 1B,C; significant main and within-regime effects of Validity on both error rate and mean correct RT (all p<0.001), and a Validity x Regime interaction in both error rate, F(4, 76) = 27.2, p = 4.1e-08, and mean correct RT, F(4, 76) = 3.23, p = 0.040, reflecting the markedly weaker biases in the Easy regime). More misses (trials with no response made within the 1600-ms motion period) occurred in the LoCoh regime than Easy and fewer occurred in the Deadline regime (main effect Regime: F(2, 38) = 28.9, p = 2.35e-08, all pairwise comparisons p<0.005; see Figure 2) and misses also scaled with validity (F(2, 38) = 3.6, p = 0.035; all pairwise comparisons p<0.05 except Invalid vs Neutral). Consistent with starting point biases being the dominant adjustment for prior probability (Mulder et al. 2012), RTs for correct trials were fastest for valid trials, whereas RTs for error trials were fastest for invalid trials, in both the Deadline and LoCoh regimes (rm ANOVAs: Validity x Correctness interaction p<0.005 in both regimes). Finally, mean RT for errors was significantly faster than correct responses in the Deadline regime, but not in the LoCoh regime which had an opposite tendency (Regime x Correctness interaction in rm ANOVA omitting Easy regime due to lack of errors: F(1,19)=59.1, p=3.03e-7; Correctness effect p=2.34e-06 in Deadline, p=0.12 in LoCoh).

**Figure 2:**
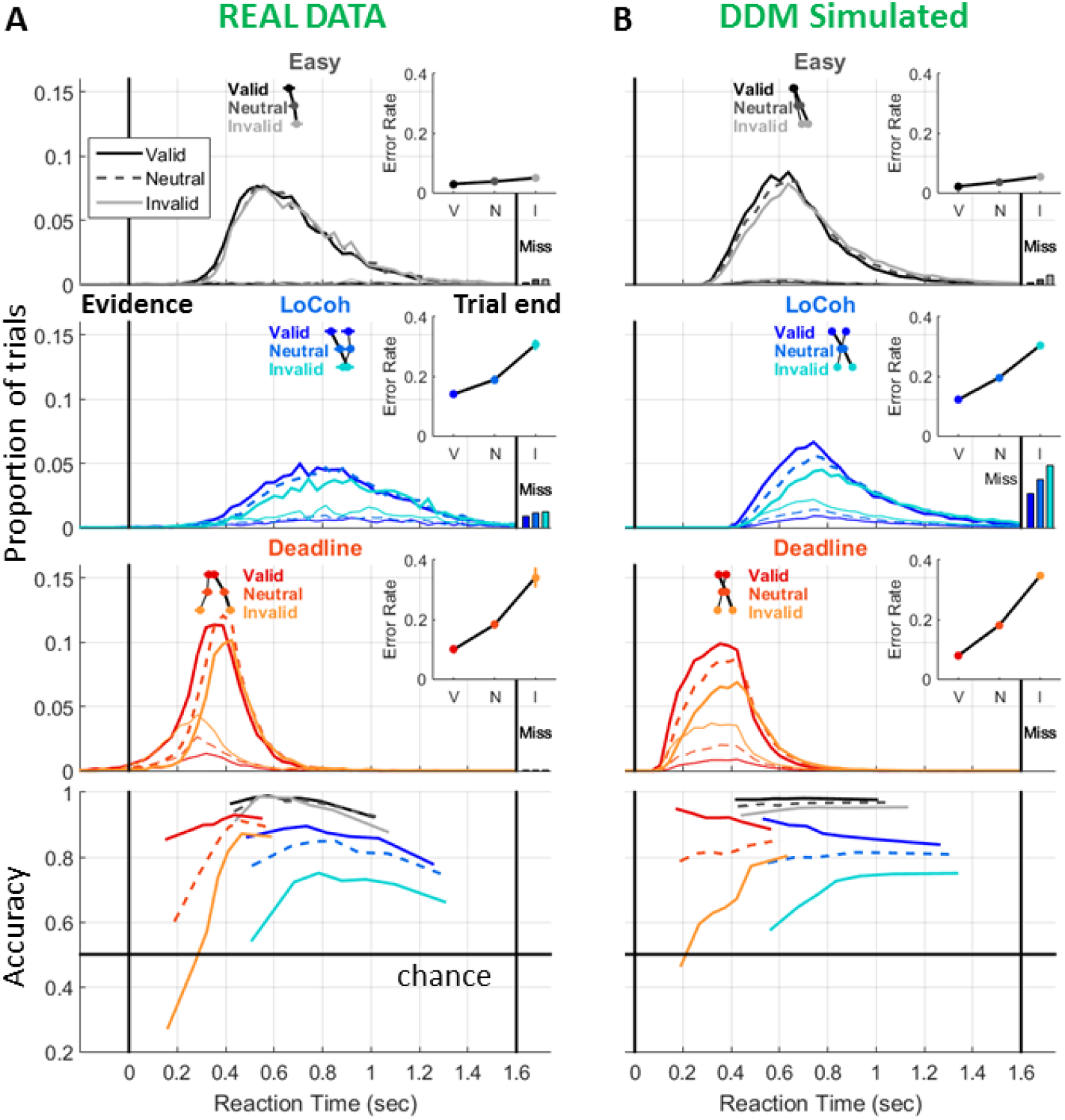
A) Empirical response time (RT) distributions for correct (thick) and incorrect (thin) responses under Easy, weak-evidence (‘LoCoh’), and time-pressured (‘Deadline’) regimes, pooled across subjects. The proportion of misses (no response within 1600 ms of evidence onset) for each condition is shown as a bar on the right. Mean RT reproduced from Fig 1B is marked above the RT distributions, and average error rate reproduced from Fig 1C are plotted in the insets, to aid comparison with model simulated data. At the bottom, conditional accuracy functions are shown for all conditions, quantifying the accuracy in each of 7 equal-sized RT bins. To follow up on a Regime x Validity x RT-bin interaction (F(24, 456) = 14.3, p = 2.22e-16), a Validity x RT-bin repeated-measures ANOVA was carried out in each Regime and revealed significant main effects of both factors in all 3 regimes (all p<0.0005), and a significant Validity x RT-bin interaction in the Deadline and LoCoh Regimes (both p<0.001, p=0.058 for Easy). Follow-up tests focused on the inverted-U nature of the functions showed that the slowest RT bin had significantly lower accuracy than the 2nd-slowest bin for LoCoh and Easy (both p<0.002; but not Deadline, which lacked slow RTs in general), and the fastest RT bin had significantly lower accuracy than the 2nd-fastest in all 3 regimes (all p<0.0001). B) Simulated behavioral data from the constant-bound, Drift Diffusion Model (DDM) allowing drift rate biases, plotted in an identical way.

### Standard diffusion model fit

To provide a more comprehensive view over the behavioral data we plotted the RT distributions for all conditions, along with conditional accuracy functions, which quantify accuracy for each of a series of equal-sized RT bins (Figure 2A). For later comparison with the neurally-informed model, we first fit a standard drift diffusion model (DDM) to these data, using a “full” version (Vandekerckhove and Tuerlinckx 2008) which included between-trial variability in non-decision time, starting point and drift rate, and also allowed mean non-decision time and drift rate to vary across regimes as well as bound separation. To capture the biases due to the prior probability cues, we included both a starting point bias and a drift rate bias parameter for each regime. The inclusion of drift rate bias was based on the measurement of significant prior probability effects on accuracy for even the very longest RTs (Paired t-tests on 7th RT bin, Easy: p=0.034139; LoCoh: p=4.7337e-05; Deadline: p=0.021933), a qualitative signature of drift rate bias that tends not to be predicted by starting point biases alone (See Figure 2 – figure supplement 1B). As expected, this DDM was able to capture all of the main effects on accuracy and mean RT across regimes, prior cue validity and correctness (Figure 2B). Further, estimated model parameters suggested interesting effects of Regime (Table 1). In addition to the expected strong decrease in decision bound under speed pressure, non-decision time was dramatically shortened by >150 ms in the Deadline regime, and was >100 ms longer for the LoCoh regime. Further, in addition to the expected reduction in drift rate due to lower physical coherence (LoCoh regime), drift rate was also lower for the Deadline regime compared to the Easy regime, despite involving the same motion coherence. These effects replicate previous behavioral modeling studies examining nondominant speed-accuracy effects (Voss et al. 2004; Rae et al. 2014; Dutilh et al. 2018), and were significant in the sense that Akaike’s Information Criterion (AIC) values were lower for this full model compared to model versions that either imposed one non-decision time across regimes or a single drift rate per coherence (Table 2). However, the improvement in fit due to including drift rate biases was very slight and did not overcome the parameter penalty of the AIC, presumably because the main identifying expression of this type of bias – accuracy differences for long RTs – represents only a small part of the behavioral data.

**Table 1:**
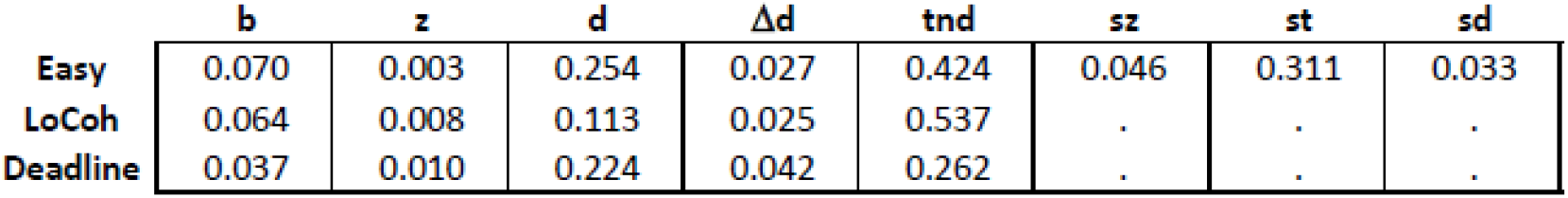
estimated parameter values for a ‘full’ diffusion model fit to the behavioral data. **b** refers to the bound, **z** the starting point bias, **d** the drift rate, **Δd** the drift rate bias (the amount added to mean drift rate **d** for valid trials and subtracted for invalid trials), **tnd** the non-decision time, **sz** the starting point variability, **st** the non-decision time variability, and **sd** the drift rate variability. Within-trial Gaussian noise is fixed at a value of 0.1, acting as the scaling parameter for the DDM. The three variability parameters each take a single value across all regimes.

**Table 2:**
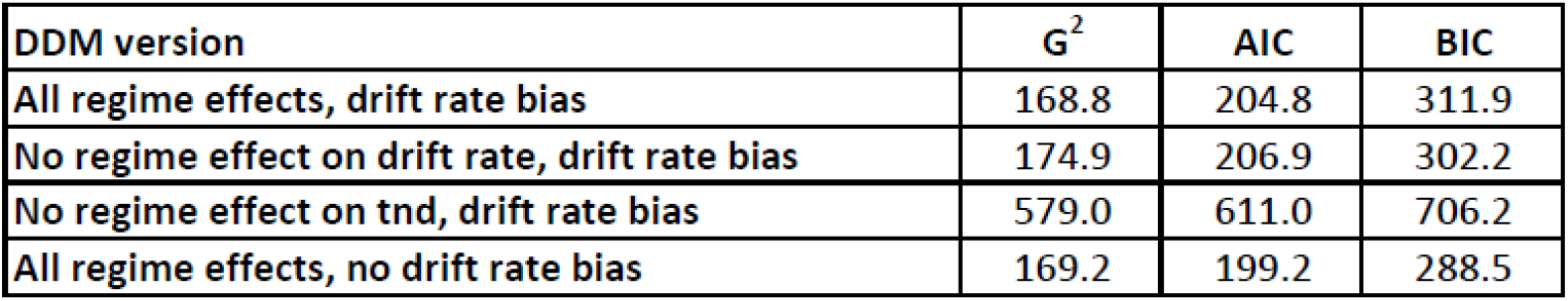
goodness-of-fit metrics for full DDM and versions of the model with each non-dominant regime or prior probability effect omitted in turn. We provide G^2^ as well as the AIC and BIC metrics which respectively penalise less and more strictly for complexity for reference. Given the purpose of the study to establish the full range of effects with convergent support from neural data, we use AIC for model selection throughout.

| DDM version | $G^2$ | AIC | BIC |
| --- | --- | --- | --- |
| All regime effects, drift rate bias | 168.8 | 204.8 | 311.9 |
| No regime effect on drift rate, drift rate bias | 174.9 | 206.9 | 302.2 |
| No regime effect on tnd, drift rate bias | 579.0 | 611.0 | 706.2 |
| All regime effects, no drift rate bias | 169.2 | 199.2 | 288.5 |

While the standard DDM provided a good account of the aggregate behavioral effects, there are prominent details of the RT distributions that are clearly not convincingly captured. First, in this task, where very fast responses are not penalised beyond the awarding of zero points if incorrect or if RT<0, and very long RTs are discouraged by a deadline, RT distributions do not have an obvious rightward skew, yet the assumption of constant bound separation in the standard DDM forces a significant skew and leads to an overestimated number of misses in the LoCoh regime (e.g. predicted 3.7% compared to actual 1.1% for neutral trials; figure 2B). Second, there are many very fast responses made in the Deadline regime that appear to be purely based on the prior cue and not on the sensory evidence, which at that time has probably not yet begun to be represented in the brain. The standard DDM is incapable of producing such “fast guesses” because by construction, threshold-crossings are only permitted to be driven by the accumulation of evidence with a stationary, positive mean and the non-decision time sets an absolute minimum on RT, so that all responses must be influenced by informative evidence to some degree. Fast guess responses are thus commonly regarded as qualitatively distinct “contaminant” responses generated by separate, non-evidence based processes and excluded or cordoned-off in a guess or lapse rate parameter in mixture models (Ratcliff and Tuerlinckx 2002; Vandekerckhove and Tuerlinckx 2007). In the neurally-informed model we next describe, we attempt, instead, to explain the full range of behaviors without recourse to trial exclusion or the random mixing of alternative decision strategies.

### Neurally-informed model construction

In our model construction approach, we struck a balance between two important factors. On one hand, constraining more model parameters to directly match certain neural measurements provides greater scope for additional free parameters, allowing the model to take on greater complexity and potentially uncover a wider range of effects. On the other hand, the accuracy of the inferences generated by such a model will be closely tied to the accuracy of the assumptions regarding correspondences between neural measurements and model parameters. We thus restricted our constraints to decision signal characteristics that, in principle, should uniquely correspond to particular model parameters, namely the motor-level decision threshold, motor-level starting points and post-decision motor execution delays. Other signal features such as buildup onset and buildup rate, when measured in trial- or subject-averaged data, in contrast, do not uniquely correspond to discrete parameter values in a one-to-one manner (Purcell and Palmeri 2017). The inclusion of two functionally distinct levels reflecting evidence accumulation (CPP) and motor preparation (mu/beta) as motivated below, provided additional empirical means of validating the model (i.e. in terms of its ability to recapitulate dynamics at both levels) while also enabling it to more accurately reflect the underlying neural decision architecture.

#### Motor-level Decision Threshold

We first sought to confirm that, as in our previous work (O’Connell et al 2012; Steinemann et al 2018), motor preparation signals reach a constant level at response, consistent with a fixed action-triggering threshold set at that level as observed in sensorimotor regions of the monkey (e.g. LIP, Roitman and Shadlen 2002; FEF, Hanes and Schall 1996). The mapping of motion direction to left and right manual responses allows the measurement of motor preparation for the two competing alternatives by comparing sensorimotor mu/beta rhythms (“MB” signal, integrated over 8-30 Hz; e.g. Donner et al 2009; de Lange et al 2013) over left- and right-hemisphere motor areas. Here we individually trace the two competing signals contralateral and ipsilateral to the correct, motion-indicated movement in parallel, representing motor preparation for the correct and wrong choice, respectively. Plotting the MB amplitude just prior to response across all conditions and for several RT bins revealed that the motor preparation signal contralateral to the button pressed reached a stereotyped threshold level regardless of the timing of the decision, the perceptual difficulty, the speed pressure, the correctness of the response, and the prior information available (Figure 3). Thus, at the motor level of our model we were able to set a constant threshold on both of two parallel buildup signals representing the two action alternatives, and set this value to 1 as our model scaling parameter (a standard practice so that all other parameters can be interpreted relative to this anchor point).

**Figure 3:**
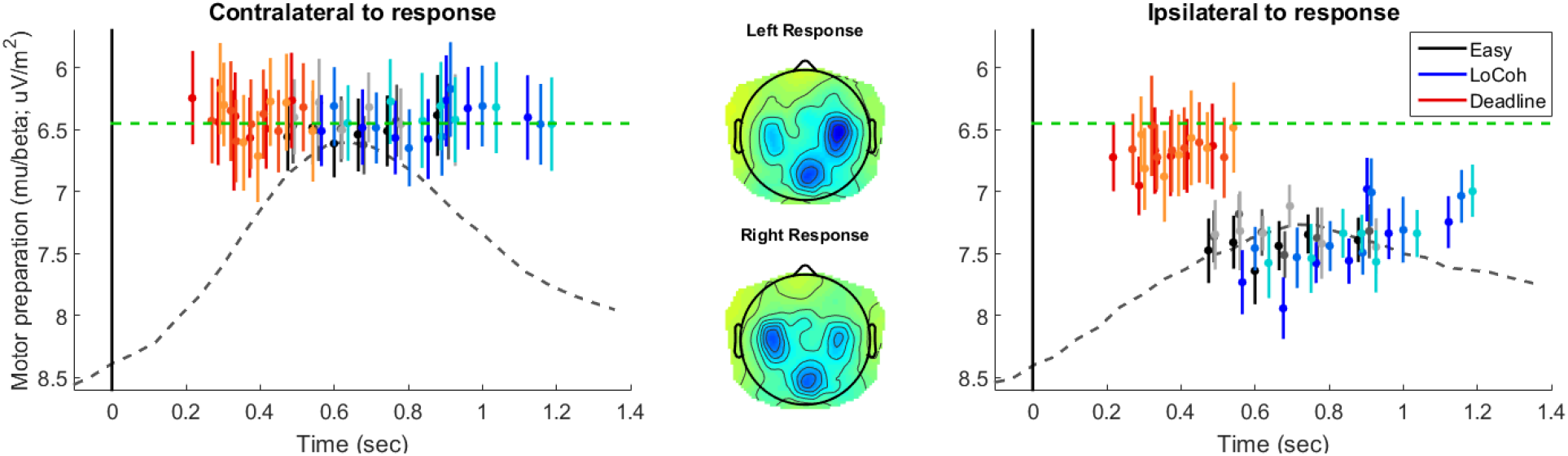
Motor-level signal for the responding hand reaches a fixed threshold level at response. Mu/Beta (MB) amplitudes contralateral (left) and ipsilateral (right) to the executed response, measured just prior to the time of response with the data for each prior condition and regime divided into 5 equally-sized RT bins for correct responses and one for errors. As a reference, the average stimulus-locked motor preparation waveforms for neutrally-cued, correct trials in the Easy regime are plotted as dashed traces in the background. Note that the y-axis is flipped so that increasing motor preparation, reflected in decreasing spectral amplitude, is upwards. Across an expansive range of RT, the contralateral MB amplitude exhibits remarkable stereotypy, indicating a fixed-threshold relationship to action execution at the motor level, whereas by comparison, MB amplitude ipsilateral to response reaches a point consistently lower than threshold and varies extensively across conditions and RT. The mean level of motor preparation at response across all conditions is indicated by a green dashed line. This is set as the threshold level in the neurally-informed model. Error bars for each point (mean ± standard error) were computed after factoring out between-subject variance.

#### Motor-level Starting-Points

Shifts in the baseline (pre-evidence) level of motor preparation have been established as the dominant neural correlate of adjustments for speed pressure and prior information (Bogacz et al. 2010; Heitz 2014; de Lange et al. 2013). We therefore next examined the neural dynamics at the motor-preparation level between the cue and evidence onset. Replicating previous studies, pre-evidence motor preparation levels were elevated under speed pressure - quite dramatically here due to the extreme deadline manipulation - and for the prior-cued response, while starting lower for the uncued response relative to neutral (Figure 4). More strikingly, in all regimes, motor preparation starts to build before sensory evidence could possibly yet impact on it. This pattern indicates the presence of evidence-independent dynamic urgency signals at the motor level in this task. A 3 x 3 repeated-measures ANOVA on the MB amplitude measured immediately prior to evidence onset (see Methods) revealed significant main effects of Regime (F(2,38)=8.08, p=0.0064) and Prior Cue (F(2,38)=6.53, p=0.0037), but no interaction between the two (p>0.1). The Regime effect was driven by the dramatic increase in motor preparation in the Deadline regime which has reached roughly two-thirds of the distance to the motor threshold with respect to the Easy and LoCoh regimes by evidence onset (p=0.0086 at 706 ms). The Cue effect reflected increased preparation of the cued response. These levels of motor preparation prior to evidence onset were set as constraints in our neurally-informed model.

**Figure 4:**
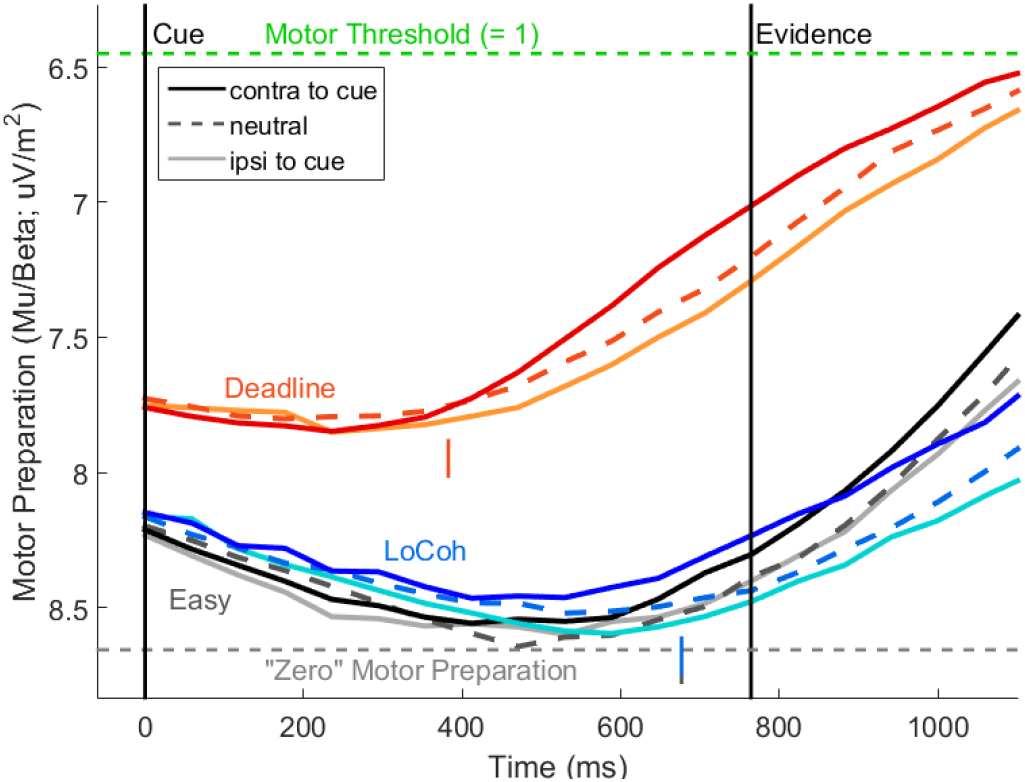
Timecourse of motor preparation (MB amplitude; 8-30 Hz) after cue onset. Vertical colored lines mark the first time point at which the temporal slope of the motor preparation signal for each regime (collapsed across cue validity) reaches a significantly positive value (382 ms post-cue for Deadline and 676 ms for both Easy and LoCoh). Note that MB is measured in approximately 300-ms windows stepped 59 ms at a time, which sets a limit on the resolution with which the time of acceleration can be precisely estimated, but is sufficient to conclude that the urgency kicks off prior to the time that evidence representations can influence motor preparation. The green ‘Motor Threshold’ marks the same level as in Fig 3, i.e., the mean level reached by MB contralateral to the response.

#### Post-decision Motor Execution Delays

We next estimated the duration of post-decision motor execution processes, regarded as part of the non-decision time construct in decision models, by measuring the onset of motor execution potentials (O’Connell et al 2012; Dmochowski and Norcia 2015; Steinemann et al 2018, see Methods). We found that execution potential onset was significantly later for LoCoh (−82 ms) than Easy (−97.5 ms), and later again for Deadline (−69 ms, both p<0.05, jackknife t-tests), but did not significantly differ across cue validity conditions (p>0.38). We thus constrained the motor time, i.e. the time delay from the threshold crossing of our winning motor preparation signal until the completion of the button click, in our model to these three values, with no variation across cue validities. Although this constrains the non-decision time at the motor-execution stage of the stimulus-response period, non-decision delays also exist at the initial stimulus onset classically associated with sensory encoding processes (Ratcliff and McKoon 2008). In our model, we reasoned that the timing of evidence representation in sensory areas would differ little across the coherences used, and so a single evidence onset time parameter was used to represent the point where the mean of the noisy evidence stepped up to a positive value. Accumulation start time, however, was assumed to be set independently from evidence onset, and strategically adaptable to task regime in accordance with recent theories based on behavioral modeling (Teichert et al. 2016). Having a separate accumulation onset time per Regime allowed the model fits to estimate differences in non-decision delays in an analogous way to the standard DDM. Like in the full DDM, non-decision time was assumed to vary randomly across trials, and this variability was here applied to both accumulation start time and the motor non-decision time.

#### Distinct Evidence Accumulation and Motor Preparation Levels

Aside from the fact that it traces cumulative evidence independent of motor requirements, there are several findings that indicate that the CPP reflects a distinct processing level intermediating between sensory encoding and motor preparation. First, the evidence-dependent build-up of the CPP has been shown to reliably precede that of motor preparation signals measured on the same timescale (Kelly & O’Connell, 2013). Second, a recent investigation found that in contrast to motor preparation signals like Mu/Beta, the CPP did not adjust its starting level in response to speed versus accuracy emphasis cues and its pre-choice amplitude declined as a function of RT (Steinemann et al. 2018). Here, accordingly, we observed no significant effect of probabilistic cue instruction nor of Regime on the CPP measured at evidence onset (both main effects of 3×2 rm ANOVA p>0.5, Figure 6) and its amplitude varied systematically as a function of RT (see below). These patterns are consistent with the application of static and dynamic urgency at the motor level but not directly at the putatively “upstream” processing level reflected by the CPP, which may instead encode a pure representation of cumulative evidence. We therefore included distinct evidence accumulation and motor preparation levels in our neurally-informed model. Note that the inclusion of these two separate levels does not entail additional parameters over typical bounded accumulation models with urgency/collapsing bounds - it simply retains a separate representation of cumulative evidence and urgency for the purpose of accounting for neural signals in addition to behavior.

**Figure 6:**
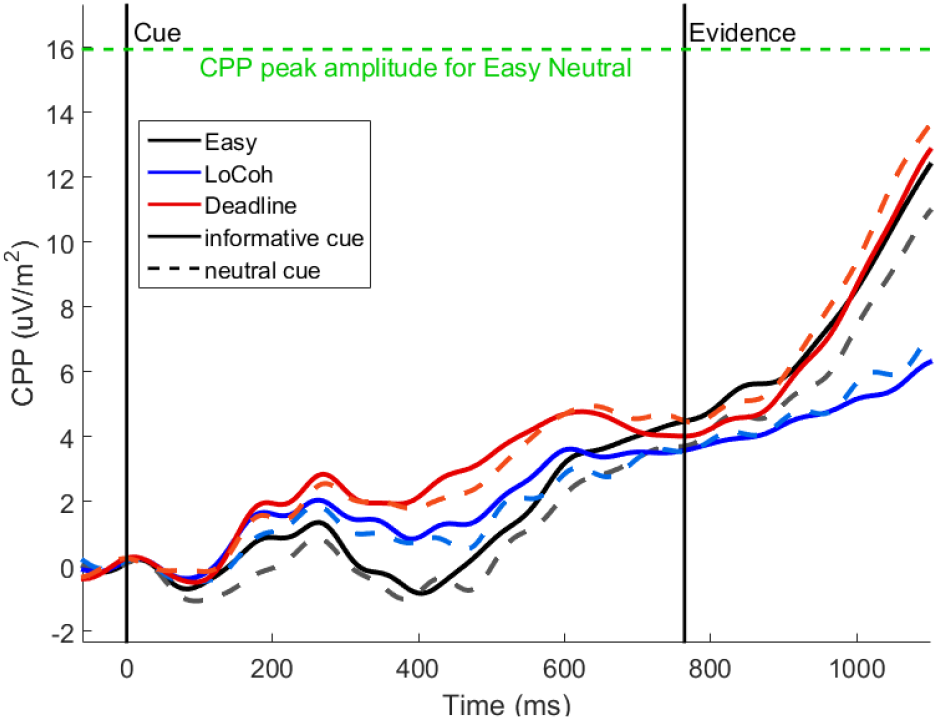
Event-related potential waveforms for neutral and informative cues in the cue-evidence period, at electrodes in the focus of the centro-parietal positivity (CPP). Note that since the color cue itself must be decoded and processed, there is a positive-going shift in the waveform during the cue-evidence interval, which is greater in the regimes that strongly call for incorporation of the prior probability information (main effect of Regime at 450 ms, F(2, 38) = 5.86, p = 0.021), but this difference dissipates by the time of evidence onset (see main text).

Thus, the above features and constraints culminated in a neurally-informed model in which noisy evidence is accumulated by an intermediate, motor-independent decision signal stage reflected in the CPP, which in turn feeds two racing build-to-threshold motor preparation signals that also receive an additive dynamic urgency signal (Figure 7). The offset of the dynamic urgency signal at the outset of the decision process is constrained by the MB amplitude values for each regime and prior cue condition, while the rate of urgency buildup is free to vary across regimes as well as randomly across trials. The intermediate accumulation (CPP) level receives no urgency input directly and represents purely the cumulative evidence bearing on the decision. Critically, the CPP is itself not directly subjected to a threshold-crossing criterion; rather, it builds as far as it can before the threshold is crossed at the motor level, such that its pre-response amplitude is nevertheless strongly affected by motor urgency. The complete model structure is illustrated in Figure 7.

**Figure 7:**
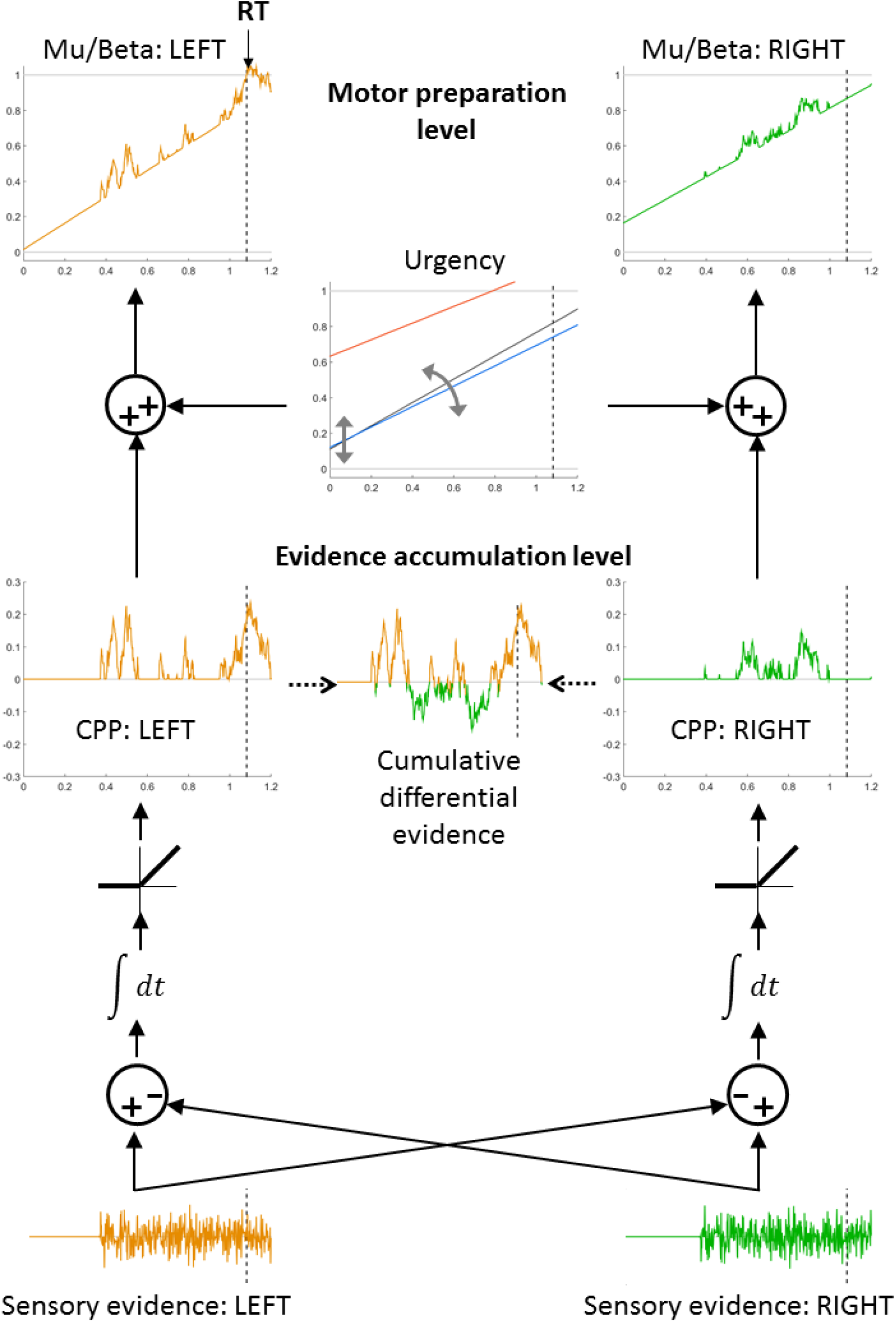
Multi-level bounded evidence accumulation model. Similar to previous models (Mazurek et al. 2003), the evidence representations for the two competing directions of motion (left and right) are subtracted and accumulated by two separate accumulators. In our model, the two competing accumulator units effectively represent the half-wave rectified cumulative differential evidence in favor of left and right respectively. This is equivalent to two positive-only populations together representing a single signed quantity of cumulative differential evidence, with one population catering for the positive and the other for the negative values. These two accumulators are assumed to sum together to produce the CPP signal that we measure on the scalp surface. Each of these accumulator units at the CPP level is fed into a motor preparation process at which an action-triggering threshold is set, and a common source of evidence-independent, dynamically growing urgency adds to both of these competing motor preparation signals. The traces in this schematic are from a single example trial in the LoCoh regime in which evidence favors the leftward direction and results in a correct response being made at 1.082 sec. We chose a trial in which the evidence was particularly uncertain to aid our explanation. The urgency signal plot shows the average urgency signal component for each of the regimes, and the gray arrows inside it signify that on each trial, the starting value and slope of the urgency signal varies randomly around that mean trajectory.

### Comparing DDM and NI model accounts

The simulated behavior from the neurally-informed model provides a compelling match to the empirical behavior, capturing not only the aggregate behavioral trends in mean RT and accuracy across regimes but also the lack of rightward skew in the distributions and biased fast guesses in the Deadline regime (compare Figure 8A and B; see Figure 8 - figure supplement 1 for model version with no drift rate biases). Accordingly, the fit quality was markedly superior to that of the standard DDM (Table 4, compared with Table 2). Table 3 lists the parameter estimates of the neurally-informed model. Several mechanistic adjustments for regime and prior probability emerge that are strikingly divergent from those implicated by conventional sequential sampling modeling both here and in the extant literature (Figure 9). Further, the model naturally accounts for key dynamic features of the neural decision signals that match predictions simulated from the NI model despite not being directly used to constrain it (Figure 10). In the following we describe each of four such key effects in turn.

**Figure 8:**
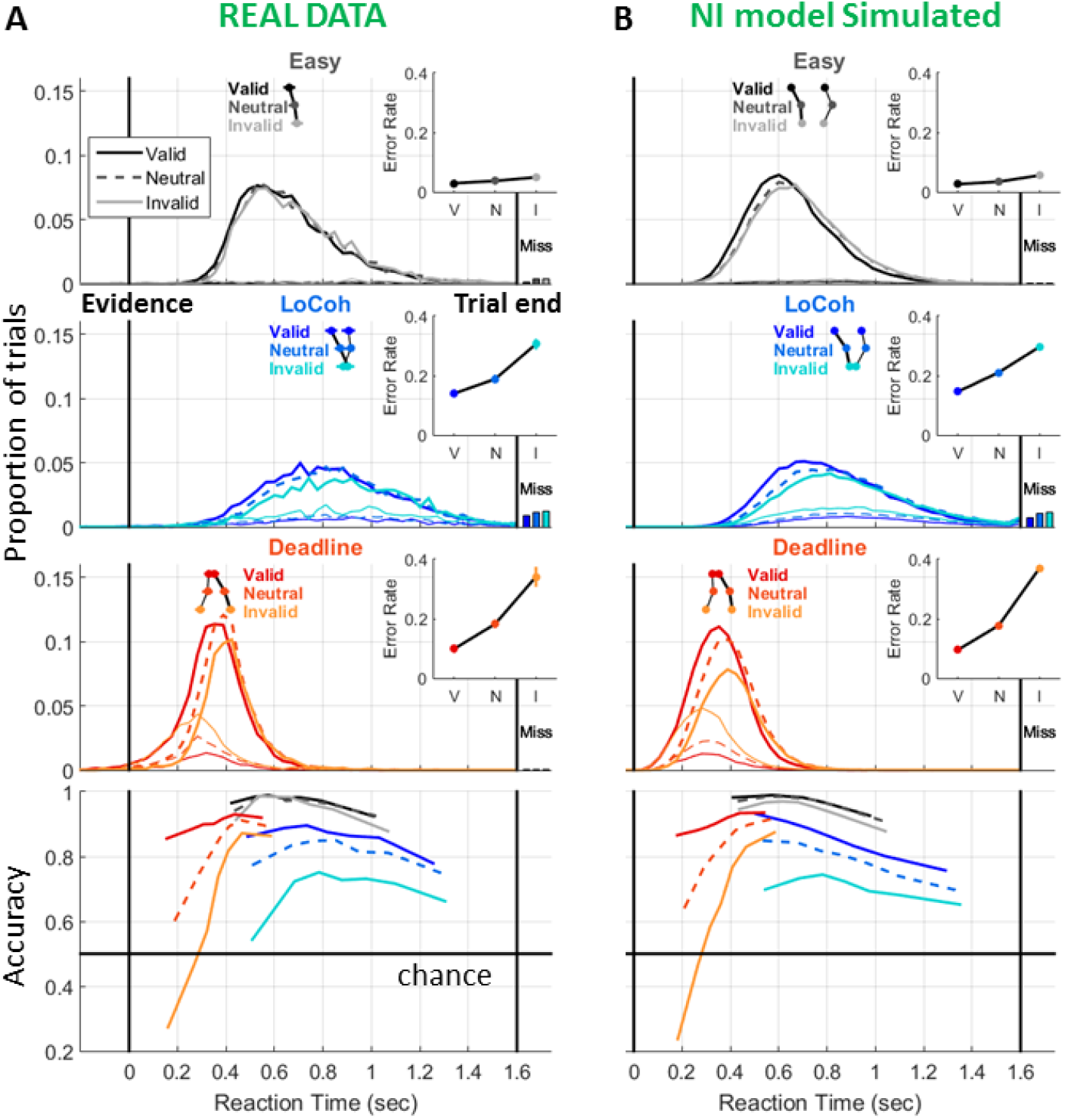
Pooled RT distributions and conditional accuracy functions, recapitulating the empirical data shown in Figure 2A (A) and this time comparing with the simulated behavioral patterns of the neurally-informed model (B).

**Figure 9:**
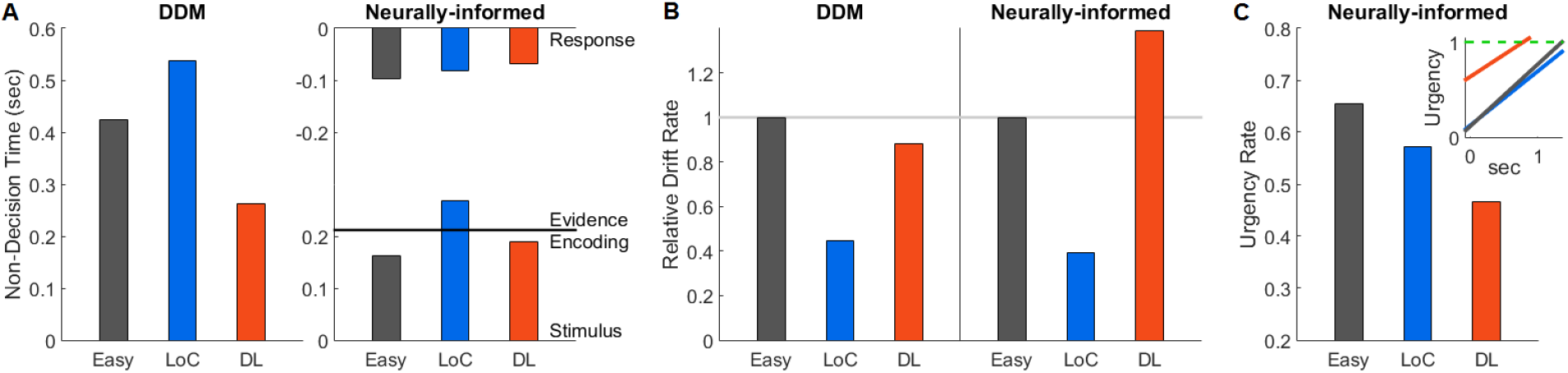
Bar plots of parameter estimates showing regime effects of the DDM and neurally-informed (NI) model. A) The DDM estimates a unitary non-decision time for each regime and suggests dramatic differences between regimes; In contrast, the NI model suggests a more subtle and distinct pattern of cross-condition effects to three separate components of the non-decision time: the post-decision motor execution delay estimated directly from neural data (shown at the top), the delay between physical presentation of the evidence and the earliest time that its representation can impact on the decision process (Evidence onset time, horizontal line), and the onset time of accumulation (lower bars). B) Estimated drift rate, normalized with respect to the value in the Easy regime to facilitate cross-model comparison. While the models agree on the marked drift rate reduction with reduced coherence, the DDM indicates that drift rate is lower under the Deadline regime but the NI model indicates that it is greater. C) Urgency rate estimated for each regime. Inset shows the linear time functions representing the full urgency signals including both offset (starting point) and slope. Although urgency has a shallower rate in the Deadline regime, it starts much closer to the threshold. Each of these three regime effects were significant in the sense that Akaike’s Information Criterion (AIC) values were lower for this full model compared to model versions that either imposed one accumulation onset time across regimes, a single drift rate per coherence, or one urgency rate across regimes (Table 4). The reliability of the regime effects in both the DDM and NI model were further verified by observing their unanimity across the 25 model fits, for which the search algorithm was started with a wide range of different initial parameter vectors, all of which were forced to not initially have any regime differences (Figure 9 - figure supplement 1).

**Figure 10:**
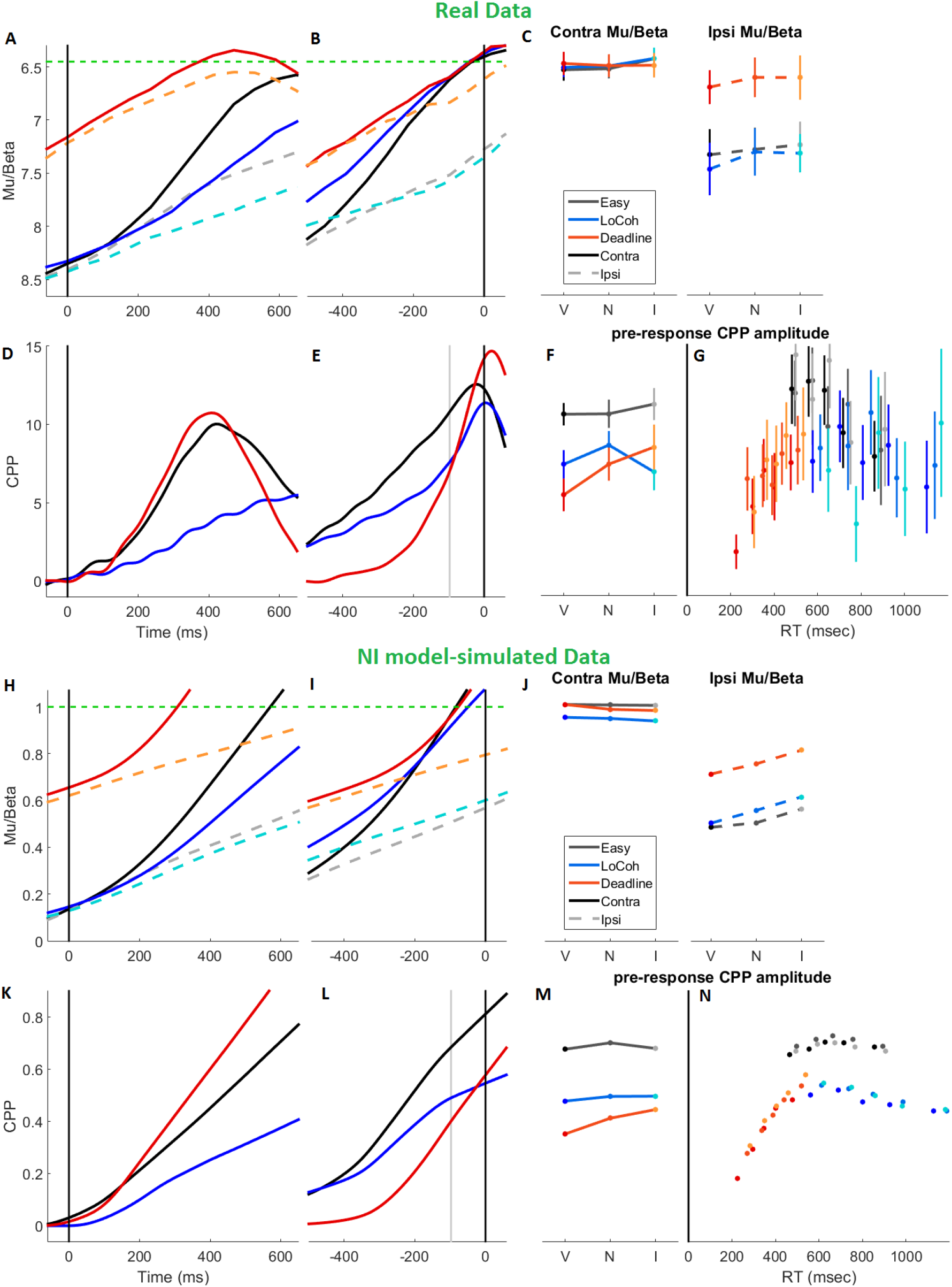
Real and simulated average decision signal dynamics and amplitudes. All data are for correct trials only. A) Stimulus-locked and B) response-locked Mu/Beta amplitude reflecting motor preparation contralateral and ipsilateral to the (correct) response executed. C) amplitudes of contralateral and ipsilateral Mu/Beta just prior to response, broken out by prior cue validity. D) Stimulus-locked and E) response-locked centro-parietal positivity (CPP) for each regime. Note that although the CPP continues to rise and peak beyond the point where we measure its pre-response amplitude (a 59-ms window centered −97.5 ms prior to response, i.e. the earliest motor execution onset), this likely represents post-decision accumulation (Resulaj et al. 2009; Murphy et al. 2015), effectively giving rise to an overshoot in the CPP’s trajectory (see Steinemann et al 2018). F) CPP amplitude broken out by prior cue validity. G) pre-response CPP amplitude measured in 5 RT bins for each prior cue type and Regime highlights an overall inverted U-shaped relationship with RT. Error bars in all panels of the real data show S.E.M. after between-subject variability has been factored out. H-N) the same plots are produced for simulated decision signal trajectories from the best fitting neurally-informed model. Note that whereas the real signals fall back or rebound at or shortly after the time of the response, our simulations continue to evolve beyond this point; because these post-decision dynamics are immaterial to the accuracy and timing of the action choices, there was no imperative to mimic these fall-down dynamics. Note also that all simulated CPP waveforms have been shifted on the time axis by a fixed amount to approximately line up with the real waveforms, in line with an estimated 134-ms delay between cumulative evidence encoding at the CPP level and its impact at the motor preparation level (see Methods).

**Table 3:** estimated parameter values for the neurally-informed model. Here **Zv, Zn** and **Zi** refer to the starting level of motor preparation relative to a threshold of 1 (acting as the scaling parameter) for valid, neutral and invalid trials, respectively, **U** refers to the urgency buildup rate, **D** the drift rate, **ΔD** the drift rate bias (the amount added to mean drift rate D for valid trials and subtracted for invalid trials), **Tac** the accumulation onset time relative to evidence presentation, **Te** the onset time of evidence encoding relative to evidence presentation, **Tm** the post-threshold motor-execution delay, **S** the within-trial Gaussian noise, **Sz** the starting point variability, **St** the non-decision time variability, and **Su** the urgency rate variability. Constrained parameter values are indicated in red, free parameters in black. Note that Te, S, Sz, St and Su each take a single value across all regimes.

| | Zv | Zn | Zi | U | D | $\Delta D$ | Tac | Te | Tm | S | Sz | St | Su |
| --- | --- | --- | --- | --- | --- | --- | --- | --- | --- | --- | --- | --- | --- |
| Easy | 0.129 | 0.073 | 0.078 | 0.656 | 1.319 | 0.077 | 0.162 | 0.212 | 0.098 | 0.522 | 0.404 | 0.225 | 0.166 |
| LoCoh | 0.156 | 0.088 | 0.056 | 0.573 | 0.520 | 0.073 | 0.269 | . | 0.082 | . | . | . | . |
| Deadline | 0.695 | 0.606 | 0.565 | 0.467 | 1.830 | 0.114 | 0.189 | . | 0.069 | . | . | . | . |

**Table 4:**
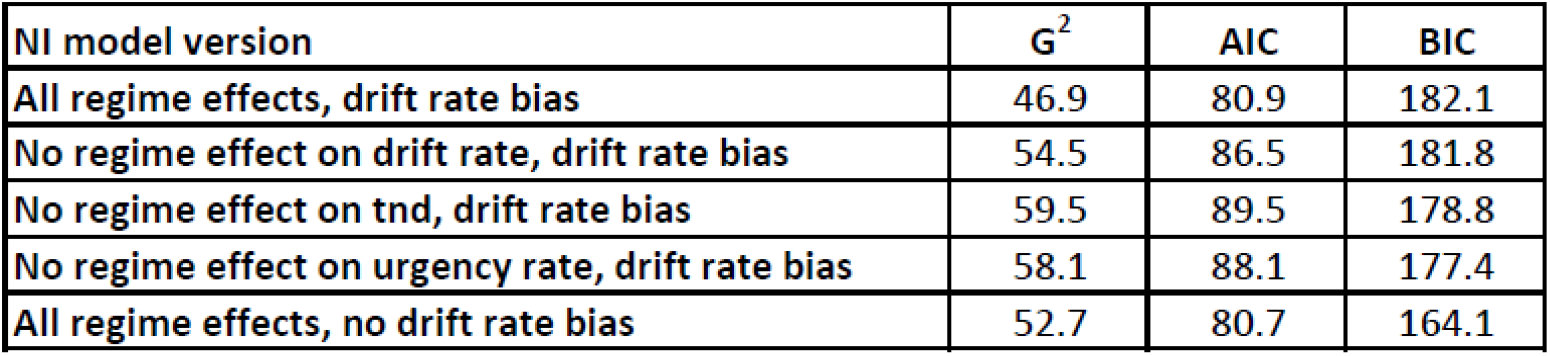
goodness-of-fit metrics for full neurally-informed model as well as versions of the model with each non-dominant regime or prior probability effect omitted in turn.

### Regime effects on non-decision time

The NI model highlighted a markedly different pattern of non-decision time effects to that identified by the DDM. Whereas the DDM estimated that non-decision time is shortened by >150 ms in the Deadline regime relative to Easy, the NI model identified differences that were an order of magnitude smaller with a small shortening (~30 ms) of motor execution times (constrained to match the neural data) offset by a small *lengthening* of accumulation onset time (~20 ms). In addition, while both the DDM and NI models indicated circa 100-ms increases in non-decision time in the LoCoh regime compared to Easy, the NI model could specifically apportion this effect to a delayed onset time of evidence accumulation as opposed to evidence encoding or motor execution delays, the latter being in fact slightly shorter. The NI model goes further to specify that the estimated onset times entail accumulation beginning before there is informative, direction-selective sensory evidence in the Easy regime (thus accumulating only noise) and afterwards in the LoCoh regime. Further, due to the combination of this pre-evidence accumulation and the early launch of the urgency signals, the NI model, unlike the DDM, was able to account for the fast biased guess responses in the Deadline condition.

To further test these model-predicted Regime effects on accumulation onset time as well as the other effects we describe next, we plotted CPP and Mu/Beta waveforms averaged across cue validities to maximize signal robustness (Figure 10A,B,D,E; see figure supplement for traces broken out by validity), and compared them with average decision signal trajectories simulated from the neurally-informed model (Figure 10H,I,K,L). Overall, the stimulus-locked waveforms around the time of accumulation onset appear well-matched across the real and simulated data. In particular, the difference in accumulation onset time between the two high-coherence regimes (Easy and Deadline), marked by the acceleration of the stimulus-locked CPP, appears negligible in contrast to the 150 ms difference estimated by the DDM (Figure 10D). To assess this quantitatively, we compared the time at which the stimulus-locked CPP in the Easy and Deadline regime reached 20% of its maximum value, and this differed by no more than 1 ms on average (jackknife t-test, p=0.95). Note that the true time of accumulation onset is not possible to precisely estimate from the CPP waveforms due to the gradual nature of initial noise accumulation, the timing jitter across trials and subjects, and the low-pass filtering of the CPP necessary to remove the SSVEP (see methods). Nevertheless, this test of the time to reach the 20% mark serves as a robust proxy and could certainly be expected to detect any large differences on the order of those estimated by the DDM. Though the differences are slight, the neural data in fact trends in the direction of evidence accumulation starting sooner in the Easy regime than the Deadline regime, consistent with the NI model parameter estimates and directly opposite to the later onset suggested by the DDM. An interesting consequence of the variable and often earlier accumulation onset with respect to evidence encoding is that the trial-averaged accumulator signal begins to climb distinctly earlier than the time at which sensory evidence causes it to accelerate, a feature that is apparent in both the simulated and real CPP waveforms (one-sample t-test on signal slope in first 118 ms post-evidence, p=0.024,no significant effects of Regime or Cue in 2×3 rm-ANOVA, p>0.1).

### Regime effects on drift rate

Whereas the DDM concludes that drift rate is reduced under the Deadline regime compared to Easy, the NI model indicates that it is markedly increased (Figure 9B). Buildup rates of the empirically-measured CPP, in fact, rank in the same way across regimes as the buildup rates of the model-simulated CPP as well as the model parameter estimates of drift rate, with the CPP in the LoCoh regime tending to build more slowly than in the Easy regime and the Deadline regime building more steeply (Stimulus-locked temporal slope in 118-ms window starting at 200 ms: F(2, 38) = 21.5, p = 5.67e-07; Response-locked slope in 118-ms window ending at −97.5 ms: F(2,38) = 3.97, p = 0.027; Relative to the Easy regime, LoCoh slope decrease reached significance only in the stimulus-locked waveforms, p=5.5e-7, and the Deadline slope increase reached significance only for the response-locked waveforms, p=0.019; Figure 10 D,E vs K,L).

### Urgency rates across regimes

The NI model indicates that there is a substantial temporally increasing urgency component acting at the motor level in all regimes. Note that only the starting levels of motor preparation are constrained by the neural data, and urgency is free to increase at any or no rate in the fit. The model also reveals a paradoxical effect whereby the rate of increase of urgency is greatest in the Easy regime and shallowest in the Deadline regime (Figure 9C). Due to the substantial headstart the urgency signal takes in the Deadline regime, however, it still reaches the threshold far earlier. Neurophysiological data show that the response-locked motor preparation buildup rates (Fig 10A,B) rank in the same way as the urgency rates of the model, with the Deadline condition exhibiting the shallowest slope (118-ms window ending at −97.5 ms, F(2,38) = 7.16, p = 0.0022, both regimes significantly different from Easy, p<0.02), and this is recapitulated in the model-simulated waveforms (Fig 10I). Since this would be partially driven by the pre-response time period often overlapping with the pre-evidence period in the Deadline regime due to very fast responses, we also tested for differences in stimulus-locked slopes between the Easy and Deadline regimes and found that the Deadline motor preparation became significantly shallower at 206 ms, even when excluding any trials with RT faster than 300 ms to avoid potential contamination by the post-response beta rebound effect (Pfurtscheller and Lopes da Silva 1999; p=0.012).

The real (Fig 10E) and simulated (Fig 10L) CPP waveforms also confirm the expectation that unlike the stereotyped threshold level reached by motor preparation, the cumulative differential evidence reaches levels that systematically vary across regimes (Figure 10F). In fact, breaking-out CPP amplitudes as a function of RT reveals that whereas its amplitude dramatically grows over the fast RTs in the Deadline regime starting from very low amplitudes, they start high and gradually decrease in amplitude over the later and longer range of RTs for the LoCoh regime (Fig 10G,N; RT-bin x Regime interaction in rm-ANOVA, F(8,152)=3.61, p=0.046), with the amplitudes lying at higher values for the mid rage of RTs in the Easy regime. This pattern is highly similar in the real and simulated data, as confirmed by the fact that correlations between the two are so high as to have a probability of 3×10^-7^ of arising by chance alone, determined through shuffling trial conditions 100,000 times. This overall inverted U-pattern arises from the disproportionately strong influence of starting point variability in the Deadline regime due to its mean starting proximity to the bound while the decline in CPP amplitude for later RTs in the LoCoh regime can be attributed to the stronger expression of the temporally increasing dynamic urgency component (Steinemann et al 2018).

### Prior probability cues bias drift rate

As expected, the addition of drift rate bias to the model allowed it to capture the full extent of the biases in accuracy for even the trials with the longest RT (compare conditional accuracy functions in figure 8B to those in figure 8 - figure supplement 1B). While in terms of AIC, the model with drift bias produced an almost identical behavioral fit to that without drift rate bias, the neurophysiological data, provided crucial additional supporting evidence. Breaking out by prior cue validity revealed an interesting pattern in CPP amplitude: A 3 x 3 repeated measures ANOVA testing for effects of Regime and Validity on CPP amplitude prior to response revealed an interaction between the two factors (F(4,76)=3.27, p=0.046). Testing each regime individually, a significant effect of prior probability was found in the Deadline regime (p=0.00085) but not in either the Easy (p=0.54) or LoCoh (p=0.52) regime, and indeed, with drift rate bias included the model-simulated amplitudes mirrored this tendency for prior biases to disproportionately affect amplitude in the Deadline regime (Fig 10M). At first blush it would seem that the biases in the starting point of motor preparation, which are almost as large for the LoCoh regime as for the Deadline regime, should cause commensurate differences in CPP amplitude because the closer to the bound the motor signals start, the less scope for accumulation there is at the CPP level. However, as illustrated in Figure 11A, due to the growing dynamic urgency component, the negative influence of a starting point increase on peri-threshold accumulator amplitude is cancelled out by the positive influence of drift bias. To look further into this, we repeated the model fits but this time including the degree of mismatch between simulated and real CPP amplitude modulations in the objective function as a penalty term. We found that the model that included drift rate bias increasingly outperformed the model with no drift rate bias as the emphasis on capturing the CPP modulations was increased (Fig 11B).

**Figure 11:**
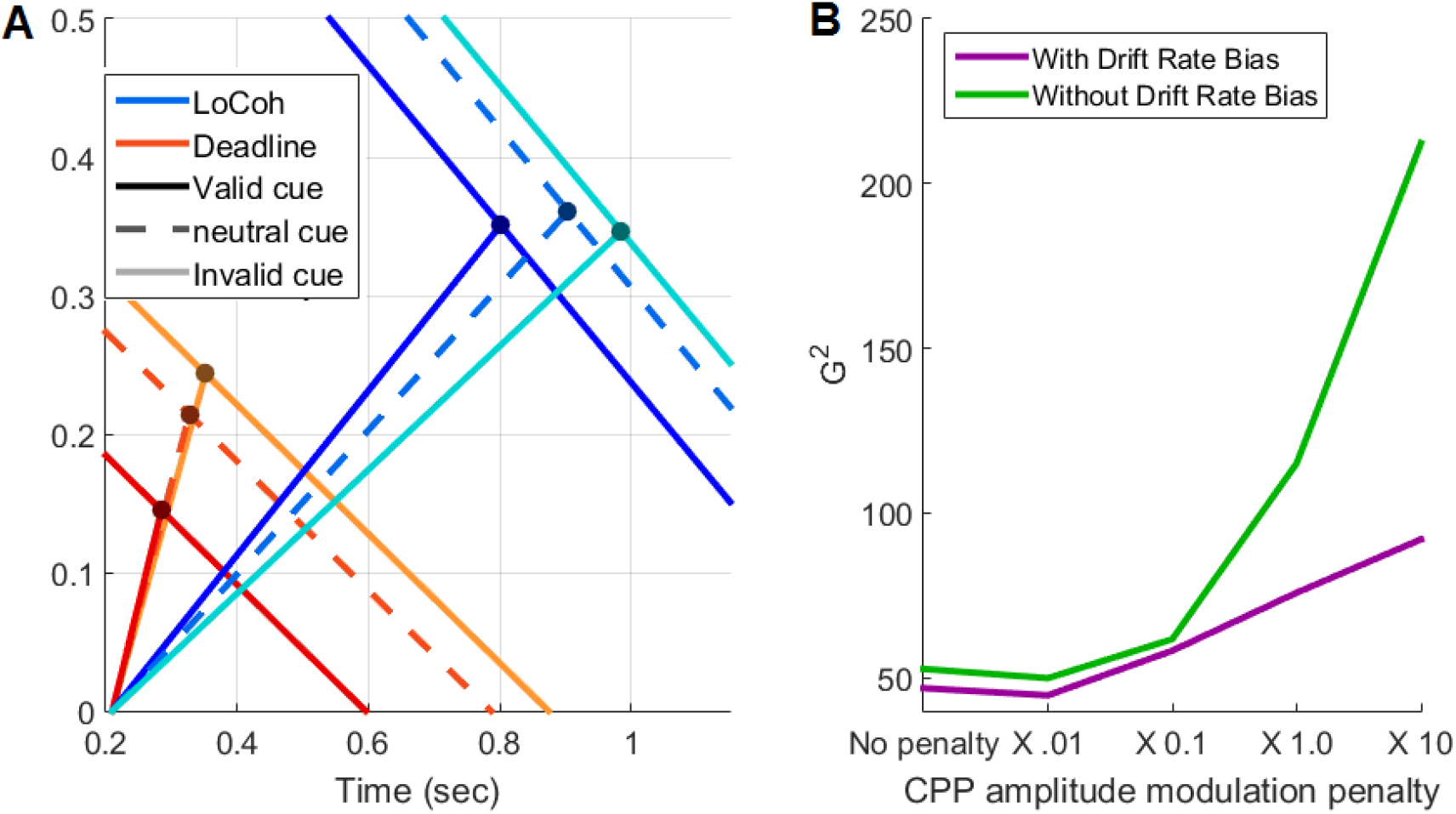
Relationship between drift rate biases and CPP amplitude. A) Predicted impact of drift rate biases on the cumulative evidence reached before the motor threshold is crossed in the LoCoh and Deadline regimes. A growing urgency signal component at the motor level effectively collapses the bound experienced by evidence accumulation, and due to prior biases also acting at the motor level, this collapsing bound is offset according to prior cue probability, as shown by the downward-sloping lines. Due to these effective bound differences, a single drift rate would result in the accumulator reaching higher levels for invalid than valid trials. However, a bias in drift rate has an influence on accumulator amplitude in the opposite direction, because the shallower the drift rate, the later and lower-down on the collapsing bound the decision variable terminates on average. In the case of the LoCoh regime in which the drift rates are all low, a bias in that drift rate leads to the drift rate bias cancelling the impact of the starting point bias on terminal accumulator amplitude, whereas for the Deadline regime which has a higher neutral drift rate, a similar degree of drift rate bias slightly opposes but does not cancel the starting point effect, and amplitude differences are still observed. The slopes and offsets of the lines in these schematics were computed directly from the model parameter values of the NI model with drift rate bias. B) G^2^ values quantifying goodness of fit to the behavioral data for model fits using an objective function with an increasing penalty term quantifying divergences of the priors effect on CPP amplitude simulated from the model from that of the real data. The larger this penalty term, the more the parameter values of the fit are dictated by the CPP amplitudes instead of the behavioral datapoints; hence, increasing this penalty term will necessarily decrease the goodness of fit to behavior. However, a greater emphasis on capturing the CPP amplitude modulations results in the model with drift rate biases becoming distinctly better than the model without drift rate biases. This reflects the fact that without drift rate biases, it is not possible to produce amplitude differences as small as those observed (as in panel A).

## Discussion

In this study we constructed a model of prior-informed perceptual decisions that can explain neural decision signal dynamics non-invasively measured from the human brain while also providing a superior quantitative account of behavior compared to the most widely-applied sequential sampling model. The constraints provided by the neural data were critical in making it possible to develop a more complex, multi-tiered model without incurring an increased number of potentially unidentifiable free parameters. While the model was constrained to reflect the observed starting and termination levels of the motor-level decision signals, it furnished novel predictions for signal dynamics at the evidence accumulation and motor levels during the period of decision formation that received independent convergent support from both behavior and neurophysiology. In so doing, we demonstrate that the parameter adjustments mediating prior-informed, task-adaptive perceptual decisions can be far more diverse than conventional behavioral modeling has suggested.

Our neurally constrained model made it possible to parse what is usually a unitary non-decision time parameter into three key components - accumulation onset, evidence encoding onset and post-commitment motor execution time. The observation that speed pressure resulted in a shortened motor time is consistent with recent studies examining speed pressure effects on electromyographic activation onset with respect to response completion (Steinemann et al. 2018; Spieser et al. 2017). We further show that even with the same deadline, motor times were shortened when greater demands were placed on perceptual evaluation due to weak evidence (LoCoh regime). The model goes yet further by revealing cross-Regime differences in accumulation onset time, lending support to the recently proposed hypothesis that it may be under strategic control (Teichert et al. 2016). Our model also highlights that these components of non-decision time can undergo distinct adjustments; for example, accumulator onset is delayed in the LoCoh regime while motor time is quickened. Further, the decoupling of accumulation onset time from the time that informative evidence is encoded provides a mechanism for anticipatory accumulation in advance of the evidence, which, as we have recently shown, can play a significant role in some circumstances such as when evidence onsets are subtle and uncertain in their timing (Devine et al. 2019).

Another key discrepancy between the neurally-informed and standard Drift-diffusion models was that the latter indicated a reduction in drift rate under speed pressure while the former indicated an enhancement. In fact, previous studies examining the impact of speed emphasis on accumulator drift rate have yielded conflicting results. Most behavioral modeling studies have reported no effects (Ratcliff and McKoon 2008) but where differences have been observed they have been reductions (Rae et al. 2014; Dutilh et al. 2018). In contrast, the two electrophysiological investigations to have examined speed pressure effects on sensory evidence representations observed enhancements that were accompanied by corresponding increases in decision signal build-up rates (Steinemann et al. 2018; Heitz and Schall 2012). Our results provide convergent modeling and neurophysiological support for the latter observation, highlighting a modulation that serves to alleviate some of the accuracy cost caused by lower bound settings. Given that a standard DDM fit to the same behavioral data suggested the opposite effect, it is likely that the discrepancy arises from differences in model structure and not cross-study differences in task demands.

As discussed earlier, another contentious issue in the previous literature has been the question of whether decision bounds are fixed or dynamic (Ratcliff et al. 2016). Where the standard DDM asserts that bounds are fixed, a central feature of our neurally-informed model was a strong, evidence-independent, temporally growing urgency component that adds to the influence of cumulative evidence on motor preparation (Churchland et al. 2008; Hanks et al. 2014). Correspondingly, and replicating our previous neurophysiological observations (Steinemann et al. 2018), we found that for the regimes with a longer deadline, the decrease in choice accuracy as a function of RT was mirrored in a progressive reduction of the level the cumulative evidence signal (CPP) reached at the time of commitment. A particularly striking expression of dynamic urgency was the extremely early launch of the motor preparation signals even before evidence onset in the Deadline condition. This is not the first account of a motor-level buildup process kicking off in advance of sensory evidence accumulation; for example, the compelled response task developed by Stanford, Salinas and colleagues was designed to invoke exactly this kind of behavior and neuronal activity in FEF exhibits corresponding dynamics (Stanford et al. 2010; Hauser et al. 2018). In fact, years previously, Coles et al (1985) proposed that response channels are activated and start heading towards criterion potentially long before bottom-up sensory evidence influences them. We further revealed interesting differences in the rate of buildup of the urgency signal across regimes, which at first glance appear counterintuitive because the shallowest urgency rate is seen in the most urgent regime. However, the head-start provided by elevated starting levels in the Deadline regime is so large that the urgency component by itself nevertheless reaches levels close to threshold long before it does in the less time-pressured regimes. A potential reason for this strategy is that the Deadline regime provided an extremely tight time window in which to respond without any external cue to mark evidence onset. Consequently, the alternative strategy of starting motor preparation at a less elevated level and setting a steeper rate of increase may require temporal estimation of both evidence onset and deadline to a degree of precision that our subjects could not achieve, an idea worth future investigation. Regardless, the current study reveals that urgency adjustments can manifest in three distinct features of motor preparatory activity: their starting levels, their time of launch and the rate at which they approach threshold. Our results thus demonstrate how vital it is to take account of each of these components when making inferences regarding an observer’s response urgency since each can undergo different, potentially opposing contextual adjustments.

Drift rate biases induced by prior stimulus probability have rarely been reported in previous behavioral modeling work, and the present study affirms that they are difficult to detect using standard models and model selection procedures. Although the accuracy biases for long RTs provide a qualitative signature of their presence, conventional model comparison procedures indicated that the additional variance in the behavioral data explained by drift rate bias was insufficient to warrant its inclusion even using the more lenient criterion of AIC. In the case of the neurally-informed model, the AIC values were highly similar with and without drift rate bias, but we were able to conclusively adjudicate between the models based on their ability to account for response-locked CPP amplitude modulations, and this analysis strongly favoured the drift bias model. Theoretical work has demonstrated that drift rate biases are in fact optimal for decisions made about stimuli whose perceptual strength is randomly mixed from trial to trial (Moran 2015), and in such situations, neural decision signals in the lateral intraparietal area (LIP) have been found to exhibit a dynamic bias component that effectively exerts a drift rate bias (Hanks et al. 2011). However, our results suggest that drift rate biases may generalise beyond such scenarios since we implemented regimes in which the physical evidence statistics are the same on every trial. It may be that even in the absence of cross-trial variability in the physical evidence, drift rate biases are necessary to take account of random variations in their representation in the brain (Moran 2015). This finding complements another recent study showing strong drift rate biases due to relative reward value in severely time-constrained sensorimotor decisions (Afacan-Seref et al. 2018), and adds to evidence of drift rate biases arising from other sources such as previous exposure in recognition memory tasks (Ratcliff and Smith 2004), biases towards ‘no’ responses (de Gee et al. 2017) and choice history (Urai et al. 2019).

A significant feature of the neurally-informed model is that it has two separate levels of processing, allowing a flexible dissociation of motor and accumulation processes (Salinas et al. 2014). One consequence is that motor plans can be started before or after an informative evidence representation is encoded depending on environmental demands, as is particularly apparent in the Deadline regime of the current paradigm. The fast guesses observed in this condition would classically be labeled as contaminant responses (e.g. due to lapsing attention or the subject opting out of stimulus evaluation) and typically excluded from analysis (Ratcliff and Tuerlinckx 2002). Our model shows that such fast guesses can in fact arise naturally from a single, stochastic decision process that has been refined to meet tight temporal demands and emphasise speed over accuracy. The two-level architecture also has the profound implication that a representation of the “hard evidence” underlying a choice can be maintained separately from the external demands placed on response expediency, a decoupling that would be vital for metacognitive processing.

In general, the present findings highlight the value of neurally-informed modeling in making inferences more definitively than is possible on the basis of behavior alone. In this dataset, for example, the conclusion regarding drift rate modulations due to speed pressure in the standard DDM analysis was directly opposite to the conclusion of the neurally-informed model, and the latter had independent verification through testing differences in CPP buildup rate. In other cases the outcome of standard behavioral model comparison was quite equivocal (e.g. regarding the role of drift rate bias) but further consideration of the neural data with reference to simulated decision signal dynamics led to a stronger conclusion. Equally, however, the findings highlight the importance of generating full cognitive models from the neural and behavioral data rather than relying on tests of trial-averaged neural data alone. In some cases, such as measurements of starting point differences across regimes and prior conditions, it is clear that one can conclude the presence of the corresponding parameter differences, but in many other cases (e.g. when inferring the cause of decision signal build-up rate modulations), it is much less feasible to make qualitative predictions without reference to model simulations (Purcell and Palmeri 2017). For example, like in our previous study (Steinemann et al. 2018) we found a CPP amplitude reduction over RT, diagnostic of a collapsing bound, in the longer deadline conditions, but the Deadline regime showed the opposite trend despite the presence of dynamic urgency in the model and observed in the Mu/Beta traces. The model simulation confirmed that this increase in CPP amplitude over RT arises even with a collapsing bound due to the large degree of starting point variability relative to the low bound.

Nevertheless, neurally-informed modeling is not to be seen as a cure-all and warrants several cautionary notes. On a general level, models such as ours are still very much process models based on mathematics quite abstracted from neural implementation, especially from the level of spiking neural networks for which there are now a growing suite of powerful models (Wang 2002; Niyogi and Wong-Lin 2013). The approach also does not alleviate concerns over researcher degrees of freedom which always warrant caution (Dutilh et al. 2018). While neural constraints provide more freedom to explore potential parameter differences that would otherwise be intractable due to lack of constraints, one still reaches a limit on what can be inferred even with the neural signals guiding the process. Finally, the neurally-informed modeling approach we applied here entailed dealing exclusively with aggregate grand-average data and not individual-level fits and analyses. Whereas standard unconstrained models such as the DDM can be fit to individuals as much as groups, using neural constraints relies heavily on the reliability of neural measures, which is not always dependable in single-subject EEG. Future studies might address this through subject selection based on EEG signal quality. These cautionary notes notwithstanding, the present findings demonstrate the value in exploiting neural signals reflecting decision formation to inform and constrain behavioral models and offer a glimpse of their potential utility in clinical research.

## Experimental Procedures

Twenty-two human subjects (8 female) aged between 20 and 32 years each participated in 3 experimental sessions, the first for psychophysical training and the remaining two for EEG recordings during task performance. Two subjects were excluded from all analyses due to excessive perspiration artifacts in at least one EEG session. All subjects had normal or corrected-to-normal vision. Informed written consent was obtained from all subjects, and all procedures were approved by the Institutional Review Board at The City College of New York.

Subjects were seated in a dark, electromagnetically shielded booth, with their heads stabilized in a chin rest with forehead support placed 57 cm away from a cathode ray tube monitor (frame rate 85 Hz, resolution 1024 × 768) with a black background. They rested their hands on the table in front of them, with the forefingers of their left and right hand resting on the left and right buttons of a symmetrically shaped computer mouse, which formed the response alternatives. Eye position was monitored continuously throughout task performance with a remote eye tracker (EyeLink 1000, SR Research, 1000Hz). A trial was initiated by the subject by simultaneously pressing both mouse buttons, provided their measured eye position was within 3 degrees of a central, white, 5 x 5-pixel fixation point. After a short delay of 50 ms, gray, incoherently moving dots appeared, changed color after 647 ms (11 flicker frames), and finally became coherent after a further 764 ms (13 flicker frames, see fig 1A). Cue colors were yellow, green and cyan, assigned to indicate a 75% prior probability of leftward motion, neutral (50/50) and 75% rightward with the color-meanings fixed for a given subject but counter-balanced across subjects. The three validity colors were each equiluminant with the initial light gray color of the dots, as verified using a photometer. The dots remained at a constant coherence level for 1600 ms before the display was extinguished for all conditions. If the subject indicated the correct decision before a deadline, by a left or right-hand button press for left or right motion, respectively, he/she earned points that translated to monetary value. Trials ended with feedback text indicating the number of points won if correct and on time, or indicating zero points and the reason if a response was “wrong” or “too slow.” Subjects were instructed to maintain fixation for the duration of each trial, until feedback was presented. Subjects performed the task under three regimes, which were run in separate blocks of trials: 1) ‘Easy,’ which had high coherence (20%) and a long deadline (1600 ms); 2) ‘LoCoh,’ which had low coherence (ranging from 5 to 12 %, individually titrated; see below), and a long deadline (1600 ms), and 3) ‘Deadline,’ which had high coherence (20%) and a short deadline (ranging from 388 to 640 ms, individually titrated).

The dot-motion stimulus was composed of an average of 118, 6 x 6 pixel dots presented within an aperture of 8 degrees diameter (density 2.3 dots/degree squared) centered on fixation, flickering on and off at a frame rate of 17 Hz (two refreshes on, three off; see also (Twomey et al. 2016)). The on-off flicker was incorporated originally to provide a steady-state visual evoked potential (SSVEP), but as in our other recent studies, we did not analyse this because it does not represent a sensory evidence signal specifically (Kelly and O’Connell 2013). During incoherent motion, dots were placed randomly and independently within the aperture whereas during coherent motion, a proportion of the dots were randomly selected on each frame to be displaced by a fixed distance of 0.353 degrees in either the leftward or rightward direction on the following frame, resulting in a motion speed of 6 degrees per second. The task was programmed using PsychToolbox (Brainard 1997) for Matlab (Mathworks, MA).

In order to practice the task and to determine individually-titrated levels of difficulty in the Deadline and LoCoh regimes, subjects first underwent a training session. In this session, a series of short blocks of 24 trials were first run with a 1600 ms deadline, starting at 70% coherence, and stepping down by 10-20% upon achieving >90% correct responses at each level, until the subject reached 20% coherence and was fully accustomed to the fixed timing of events and the motion speed. A 60-trial adaptive staircase procedure (one-up, three-down, (Wetherill and Levitt 1965) was then run to estimate the coherence level at which the subject performed at 80% accuracy with the same 1600-ms deadline, and this was repeated until a stable level was indicated. Next, the deadline was changed from 1600 ms to the median RT from the last 20%-coherence block and 48 trials were run to estimate the subject’s typical RT distribution under time pressure. The deadline was then further shortened toward the median RT of that fast block to further increase time pressure, and provided that participants reported experiencing intense time pressure, this value was set as the deadline in the Deadline condition. As a final step in the training session, the color cues were introduced. The subject performed a practice block of 96 trials of each of the three finalized task conditions to end the training session.

Over two EEG recording sessions following the training, each subject performed 10 blocks of 96 trials of each of the three regimes (two subjects spread the 30 blocks over 3 sessions due to fatigue). Five blocks of a given regime were run consecutively at a time to ensure that subjects would settle into a strategy and to minimize cross-over. The orders of the regimes were randomized across sessions and subjects. Small adjustments to the LoCoh coherence level or to the tight deadline in the Deadline condition were made by the experimenter as necessary from block to block when the difficulty appeared too easy or difficult.

To minimize the degree to which blocks of the different conditions differed in overall value, subjects were rewarded 40 points per correct trial in the Easy condition, 50 points in the LoCoh condition (recall titration for 80% accuracy) and 70 points for the Deadline condition (less than double the easy condition to allow for further reduction in median correct RT with practice).

Using this scheme, across the twenty subjects, the average reward per trial turned out to be 40.6, 40.4 and 45.5 points for the Easy, LoCoh and Deadline conditions, respectively, which, though small, was a significant difference (ANOVA F(2, 38) = 8.43, p = 0.007, post-hoc differences significant for Deadline compared to both Easy and LoCoh (both p<0.01)). One point corresponded to $0.0031, and at the end of the experiment two blocks were randomly selected within each regime, and the total points of those 6 blocks translated into a dollar amount in compensation for the subjects’ time, in addition to $12 per hour for training and EEG setup time.

### Behavioral data analysis

We measured response times (RT) relative to the onset of coherent motion evidence. The first button click following the prior cue was always logged as the response for a given trial even if made prior to evidence onset. Such negative RTs were regarded as a legitimate manifestation of the mechanisms employed to perform the task, rather than as contaminant responses (Ratcliff and Tuerlinckx 2002). Similarly, responses occurring after the deadline in the Deadline condition were included in all analyses and modeling. Trials on which no response was made by 1600 ms were counted as misses. These trials are shown in the behavioral data (Figure 2A) but excluded from electrophysiological analysis because they cannot be response-locked. To compute RT distributions for plotting (Figure 2A), we divided trials into non-overlapping RT bins of width 35.3 ms starting from −176 ms and ending at 1600 ms) and separately counted correct trials and error responses falling into each RT. This was done for each individual subject, expressed as a proportion of the total trials for that subject, then averaged across subjects. To compute conditional accuracy functions (CAF), we first divided each subject’s pooled (correct and error) RT distribution into 7 equal sized bins, then computed the proportion of correct trials within each of these RT bins and finally plotted response accuracy over the bins’ mean RTs for each condition, averaged over subjects (Figure 2A).

### Electrophysiological data processing and analysis

Continuous electrophysiological data were recorded in DC mode from 97 scalp electrodes with a sampling rate of 500 Hz and an online reference at site FCz (ActiCap, Brain Products). All offline analysis was performed using in-house Matlab scripts (MathWorks, Natick, MA) with raw data-reading, channel interpolation and topographic plot functions from the EEGLAB toolbox (Delorme). In offline analysis, continuous data were first low-pass filtered by convolution with a 58-tap hanning-windowed sinc function designed to provide a 3-dB corner frequency of 41.5 Hz and a local extremum of attenuation coinciding with the mains frequency (60 Hz), while also avoiding phase distortion and ringing artifacts (Widmann and Schröger 2012). The data were further high-pass filtered with a cutoff of 0.1 Hz to remove slow drift (3rd-order Butterworth). Data were epoched from −176 ms to 2600 ms relative to prior cue onset (color change) and baseline-corrected with respect to a 118-ms interval centered on t = 0 ms (2 cycles of the SSVEP). Channels with excessively high variance with respect to neighboring channels and channels that saturated or flat-lined during a given block were identified and interpolated (spherical splines). The single-trial EEG data were then transformed to current source density (Kayser and Tenke 2006) to reduce the spatial blurring effects of volume conduction, in particular reducing the overlap of frontocentral and occipital negativities with centroparietal electrodes where the CPP is measured (Kelly and O’Connell 2013; Twomey et al. 2015). Trials were rejected from analysis if a blink was identified in the eye tracker trace, or if the maximum absolute CSD-transformed amplitude exceeded 600 uV/m2 for any electrode, at any time up to 150 ms following the response. This resulted in 3-29% (mean 11%) of trials being rejected.

Cue-locked (−176 to 1100 ms with respect to the color-change prior cue), evidence-locked (−59 to 650 ms with respect to coherence onset) and response-locked (−500 to 59 ms with respect to the button press) ERPs were extracted from the longer single-trial epochs. Evidence-locked and response-locked waveforms were baseline-corrected with respect to the 59-ms interval just prior to evidence onset. An additional offline low-pass filter (103-tap windowed sinc with 3dB attenuation at 6 Hz and 90 dB at 17 Hz) was applied to the ERP traces to remove SSVEP oscillations prior to plotting. Mu/Beta-band activity was measured using a short-time Fourier transform (STFT) applied to windows of 294 ms (5 SSVEP cycles) stepped by 59 ms (one SSVEP cycle) at a time, and by taking the mean amplitude in the range 8-30 Hz but omitting the 17-Hz component to avoid contamination by the SSVEP. Mu/Beta motor preparation indices were measured at standard 10-20 sites C3 and C4 (Gratton et al. 1988), while the CPP was measured as the mean amplitude in a cluster of 4 electrodes between standard sites Pz and CPz. The topographic foci were identified based on response-locked signal topographies that were collapsed across all conditions and thus impartial to the effects being tested for in the study.

For the analysis of motor threshold level in Figure 3, Mu/Beta (MB) amplitude was measured at the first STFT timepoint following −97.5 ms (the earliest motor execution potential onset in the Easy regime, see below) with respect to the button click time. To make the scatter plot we took each of the 9 task conditions (regimes x prior cues) and after shaving off the outlying 5% on each side of the distribution, divided the distribution of MB amplitude values into 5 equal sized bins and plotted the mean amplitude against the mean RT for each bin. We also overlaid the averages for error trials without dividing into quantiles due to low trial count. We measured motor-level starting points for statistical testing and for model constraints using the last STFT point prior to evidence onset, 706 ms post-cue. For Figure 4 we carried out running t-tests against zero for the temporal slope quantified as simply the consecutive two-point difference of the MB motor preparation signals and marked the timepoint at which this became significantly positive and remained so thereafter.

Since MB signals represent spectral amplitudes estimated in approximately 300-ms intervals, the temporal resolution is far too low to examine what may be subtle differences in the post-decision non-decision time. We therefore traced the higher-resolution timecourse of broadband event-related potentials measured at the same standard motor sites (C3/C4), where, just before responses have been fully executed, a sudden positive-going deflection marks the initiation of cortical processes generating the action (Figure 5; (Vidal et al. 2003)). To ensure the suddenness of this potential is conveyed clearly we did not apply the additional low-pass filter for this plot. In keeping with previous studies (O’Connell et al 2012; Dmochowski et al 2015; Steinemann et al 2018), we take the inflection point of this signal as the point of commitment to a decision alternative and onset of motor execution. To accurately estimate these inflection points we computed the first timepoint at which the first derivative (slope) of the signal shifts from negative to positive, after applying the low-pass filter to reduce the impact of noise. (Figure 5 - figure supplement 1). To test for significant differences in this marker of the onset of motor execution (Figure 5), we carried out a jackknifing procedure where the zero-crossing time, relative to the neutral Easy condition as a reference, was computed for the average signals of 19 subjects at a time, with each subject systematically excluded in turn, and scaled the standard errors of the estimates accordingly in t-tests of the differences (see (Miller et al. 2009)).

**Figure 5:**
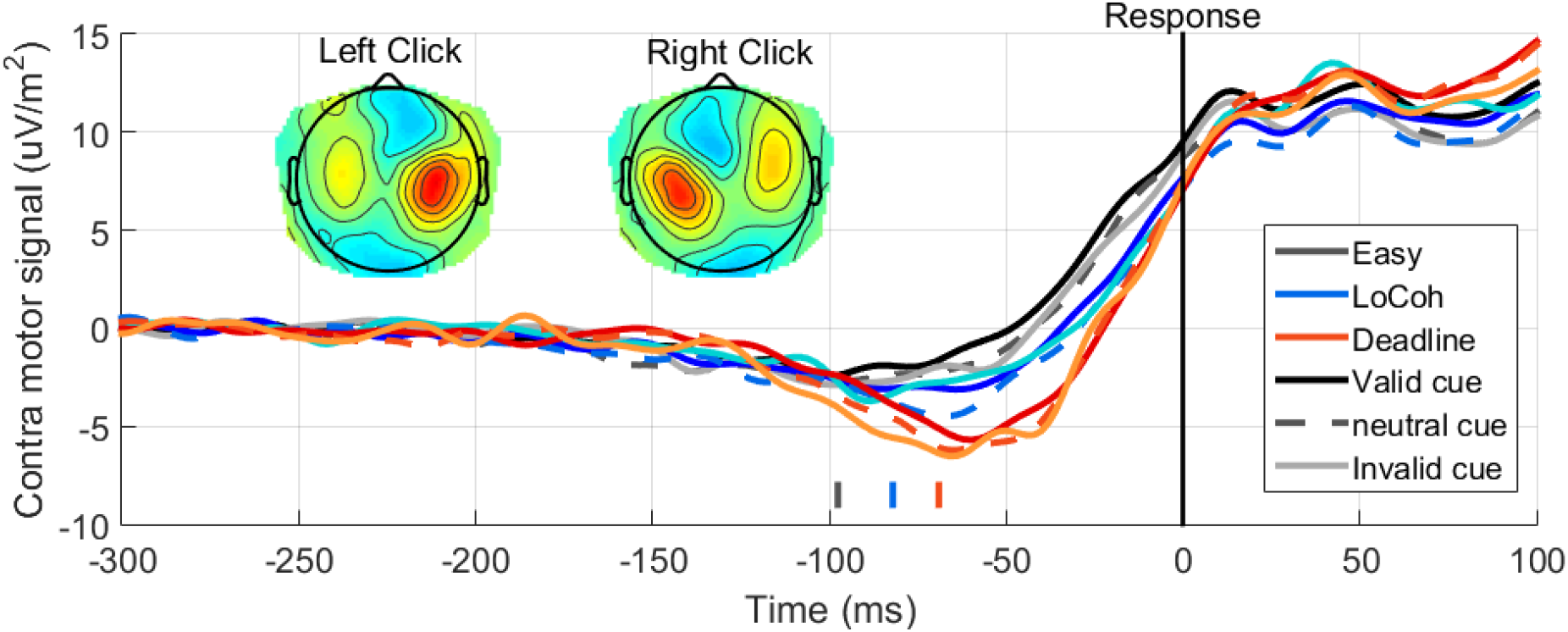
Contralateral, response-locked, motor-evoked potentials (C3 for right button presses averaged with C4 for left button presses) for correct trials. The time point at which the first-derivative signal for each condition passes through zero, indicated below the waveforms, marks the inflection point for the motor signal, where negative-going motor preparation activity gives way to a sharp potential assumed to reflect motor execution.

Pre-evidence CPP amplitude with respect to a pre-cue baseline, used to test for starting point effects of regime and prior cue validity, was measured by integrating across a 59-ms window ending at evidence onset. Pre-response CPP amplitude (Figure 10) was measured in a 59-ms time window centered on −97.5 ms, the earliest inflection point of the contralateral motor cortical ERP (C3/C4) estimated as above. CPP signal slope was tested for early accumulation by fitting a line to the 118-ms period beginning at evidence onset. CPP temporal slope differences across regimes were also tested by again fitting a line in 118-ms windows, starting from 200 ms post-evidence and ending at −97.5 ms pre-response. The 118 ms (2 SSVEP cycles) window length was chosen as a compromise between the increased robustness of measurement bought by longer time windows and the need to ensure the window does not extend beyond the period of evidence accumulation for most trials, which is very short for the Deadline condition. The equivalent was carried out for Mu/Beta by taking the temporal difference between two consecutive STFT data points together covering a 118-ms period.

### Statistical analyses

Repeated-measures t-tests and ANOVAs were employed as appropriate to test for differences in behavioral and neural amplitude and slope measures across Regimes, prior probability conditions and outcomes. For all error bars, between-subject variability has been factored out so that only variance relevant to a repeated-measures experimental design remains. For tests of signal latency, we used a jackknife procedure where times were estimated from subpopulations of subjects leaving each one out in turn, and scaling the standard error in the t-tests as appropriate (Miller et al. 2009).

### Model fitting

Models were fit to behavioral data by minimising the chi-square-based statistic G^2 quantifying the divergence of model-simulated from real behavioral data grand-averaged across subjects. To summarise the real data, we divided each of 18 RT distributions (3 regimes x correct/error x 3 validities) into 6 RT bins separated by the quantiles [.1 .3 .5 .7 .9] and counted trials in each of these bins as well as the misses for each condition, expressed these counts as proportions of overall trial counts for each subject, and then averaged both the proportions and the quantile values across subjects. Those same quantiles were then used to obtain the corresponding proportions of trials in the simulated data for any given set of model parameters. The G^2^ statistic was computed by Monte-Carlo simulation of a large number of trials (see below) and summing the divergences in trial proportions between the real and simulated data using the equation

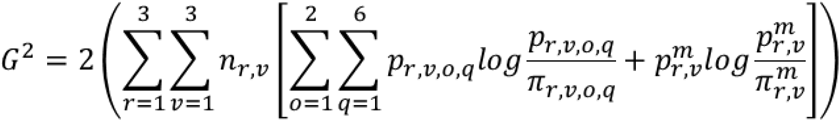

where p_r,v,o,q_ and π_r,v,o,q_ are the observed and predicted proportions of responses in bin q of outcome o (correct/error) of regime r (Easy/LoCoh/Deadline) and validity v (Valid/neutral/invalid), respectively. nr,v is the number of valid trials per regime and validity. Models with a lower G^2^ value fit the data more accurately.

We first fit a standard drift diffusion model (DDM) to the behavioral data. In its fullest form the model included 18 free parameters: a separate bound **b**, drift rate **d**, drift rate bias **Δd**, nondecision time **tnd**, and starting point bias **z** for each of the three Regimes, and a single starting point variability (uniform distribution, range **sz**), nondecision time variability (uniform, range **st**) and drift rate variability (Gaussian; standard deviation **sd**) parameter fixed across all Regimes. This model corresponds to the “full” diffusion model in common use (Dutilh et al. 2018), and allows for additional effects of Regime aside from the dominant bound adjustment, such as the modulation of drift rate and/or nondecision time (Voss et al. 2004; Rae et al. 2014), and allows for drift rate biases in addition to starting point biases (Dunovan et al. 2014). To assess the degree to which allowing for these non-standard parameter differences were warranted to optimise the fit, we also fit three other versions of the diffusion model, one that differed from the fullest model only in disallowing any drift rate biases (15 free parameters), one that differed only in setting one drift rate per coherence (resulting in 2 rather than 3 drift rates across regimes; 17 free parameters), and one that differed only in allowing one non-decision time to cover all regimes (16 free parameters). As is typical, we set the within-trial Gaussian noise standard deviation of s=0.1 as the scaling parameter.

To guide the construction of the neurally-informed model, we first focused our analyses on the decision signal dynamics at the outset of the decision process (just prior to evidence onset) with respect to the levels reached at its termination. In motor preparation signals reflected in Mu/Beta, we observed a stereotyped threshold level just prior to response and therefore constrained the motor preparation values across the nine conditions for the starting level at −59 ms relative to evidence onset. For convenience and intuitive interpretation without loss of generality, we scaled all Mu/Beta amplitude values linearly such that the average level reached just prior (first STFT point following −97.5 ms) to the response for the signal contralateral to the response corresponded to a “bound” level of 1 and the highest MB amplitude (lowest motor preparation level) during the cue-evidence interval in the Neutral Easy condition corresponded to zero motor preparation. A starting level in another regime of, say, 0.5 can thus be readily interpreted as motor preparation half-way towards execution threshold. In order to further constrain the “motor time” delay between post-threshold initiation of the cortical response execution process and the registration of the button click, we computed the time at which the first derivative (temporal slope) of the “M1” ERP signal over motor cortex contralateral to the button pressed passed through zero, indicating the inflection point at which motor execution is assumed to begin. The motor time in the model was fixed to this time point for each of the three regimes. Thus, there were 12 fully constrained parameters in the neurally-informed model. With the scaling parameter set by the bound equaling 1, we then added 17 free parameters to make the fullest neurally-informed model, including an urgency rate **U** (temporal slope of motor level urgency), drift rate **D** (mean of the Gaussian evidence distribution), accumulation onset time **Tac** and drift rate bias **ΔD** for each regime, and in common across regimes, a single parameter for each of the factors of within trial evidence noise **s**, urgency rate variability **Su**, motor starting point variability **Sz**, non-decision time variability **St**, and evidence onset time **Te**. Accumulation start time was assumed to be set independently from evidence onset, based on the idea that sensory areas represent and thus make available an evidence representation in a highly stereotyped, invariant manner from trial to trial, but the time at which that representation begins to be accumulated can vary both randomly and as a function of task demands (Teichert et al. 2016). Non-decision time variability was equally apportioned to the accumulation start time and the motor non-decision time as it was deemed likely that having separate timing variability parameters would lead to a lack of constraints. Thus, like the fullest DDM, the fullest NI model allowed for Regime effects on drift rate and non-decision time (here specifically accumulation onset) and biases in drift rate, in addition to the usual dominant adjustments. However, a strong distinction is that the dominant adjustments are not free parameters - they are fixed by the motor preparation signal data. As in the DDM, we tested for the significant contributions of each non-dominant regime and priors effect by comparing the fit quality of versions of the model that, respectively, had all of the same parameters except 1) with no drift rate biases allowed, 2) with only one urgency rate fixed across regimes, 3) with only one drift rate per coherence, and 4) with only one accumulation onset time.

The fitting procedure for both the DDM and NI models followed the same general approach consisting of the following steps, each using a bounded SIMPLEX algorithm (fminsearchbnd in Matlab, Nelder Mead) for searching the parameter space: 1) we first fit a version of the model with all regime effects allowed but no drift bias, from a large number of starting parameter vectors that were initialised with no regime differences, using 10,000 simulated trials per condition; 2) starting from the parameter vectors estimated from the 25 best fits (lowest G^2^) of the first step, we refined the fit of the same model by lowering the tolerance criteria for termination/convergence and simulating more trials per condition (20,000). This step resulted in an improvement in G^2^ of 3-6% (reduction) with respect to the best fit of the first search; 3) starting from the estimated parameter vectors from those 25 fits, we fit the fullest version of the model with drift rate biases free to vary from zero. For both models, we chose an initial level of drift rate bias (the same for all regimes) by increasing it and simulating the conditional accuracy function until a similar sized accuracy difference between valid and invalid trials was produced for the slowest RT bin of the simulated LoCoh condition; 4) starting from those fitted parameter vectors, we tested for significant contributions of the non-dominant effects as described above, by fitting versions of the model with each individual effect nullified by reducing the number of parameters. For example, to test for the significance of non-decision time differences across regimes we used a model version with only one non-decision time fixed across all regimes, and the starting parameter vector used an initial non-decision time computed from averaging the best fitting parameter values across the three regimes of the previous fit. In step 1, we selected 50 initial parameter vectors at random for the NI model, but for the DDM, to go to extra lengths to ensure the wider parameter space was comprehensively explored, we first evaluated G^2^ for simulations of 1,000 trials per condition for a coarse grid of 668,250 parameter vectors widely varying bound, drift rate, nondecision time and all three inter-trial variability parameters but keeping all parameters fixed across regimes and starting point and drift rate biases set to zero. For each combination of the three variability parameters in this set, we took the [b, d, tnd] combination giving the lowest G^2^ value and started the first coarse search from these 405 initial parameter vectors. Thus the best 25 fits in the first step were taken from 405 preliminary fits in the case of DDM, and from 50 in the NI model. The use of a comprehensive grid search to initially identify viable starting parameter vectors, which was done for the DDM, was not necessary to carry out for the neurally-informed model fit because the neural data provided a natural guide for parameter ranges in which to randomly choose values. In the case of the NI model fit, this second refinement step additionally involved expanding from 1 accumulation onset time to 3, because it was not certain whether additional regime differences in this would be needed given that motor times were constrained. To facilitate convergence of the SIMPLEX algorithm, the same random seed was used for all runs of the simulation function for each of the steps. In a very final step for both the DDM and NI models, we computed G^2^ values from the parameter estimates of each of the 25 model fits using 100,000 simulated trials per condition and a new random seed, and identified the parameter vector giving the lowest G^2^ for each of the models. To guide model comparison we computed Akaike’s (AIC) and Bayes Information Criterion (BIC) from these minimal G^2^ values, which penalise for complexity as follows,

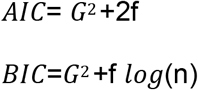

where f is the number of free parameters and n is the mean total number of trials across subjects. Simulated RT distributions in Figure 2B and Figure 8B are generated from these same simulations from the best fitting models. Although we list both AIC and BIC for reference in all fits (Tables 3 and 4), we based our model selections on AIC, because BIC penalises more severely for complexity and as a result tends to favour simple parameterizations at the expense of missing less dominant but reliable effects (Burnham & Anderson, 2004), which would run counter to the purposes of this neurally-informed study of such non-dominant effects.

To simulate the dynamics of decision formation at both the CPP and motor preparation level we simply recorded the single-trial decision variable activity timecourses at the CPP and Mu/Beta levels for each trial and averaged them in the same way as the real ERP data. To ensure consistency we filtered the noisy simulated signals using the same low pass filter in the case of the CPP and by convolving the Mu/Beta signal with a boxcar function of the same duration as the FFT window used in the real data. The CPP was simulated as the absolute value of the differential cumulative evidence, as previously done (Afacan-Seref et al. 2018). To briefly recapitulate, this is based on the assumption that the CPP reflects the activity of two neural populations that together encode a single, signed quantity of cumulative differential evidence (see Figure 7), one population representing positive values and the other negative values, though both do so with positive polarity. This accords with the observations that the CPP builds with positive polarity regardless of choice, and the fact that, despite this assumed summation of two accumulator processes, the CPP reaches a stereotyped threshold level at response for urgency-free, continuous monitoring decisions (Kelly and O’Connell 2013).

Note that without loss of generality, the model was implemented for behavior-fitting with no assumed delay between the encoding of evidence increments at the CPP level and their impact on the motor preparation level, because it is not possible to estimate this delay from behavior alone. The timing estimates related to evidence onset time and accumulation onset time should thus be interpreted as the times at which these events register at the level of motor preparation, with the understanding that their upstream representation had been encoded some time before that. For the purposes of qualitative comparison between the real and simulated CPP data, we made a rough estimate of the CPP-Mu/Beta transmission delay by simply observing differences in the timing of cross-over landmarks in the real and simulated data.

There were 4 such discernible landmarks: where the stimulus-locked Deadline regime trace crosses over the LoCoh and the Easy regimes, respectively, and where the response-locked Easy and Deadline traces respectively cross over the LoCoh trace. The average temporal shift between real and simulated data was 134 ms, and therefore we shifted the CPP waveforms leftwards by this amount of time in Figure 10. We emphasise that our purpose was not to pinpoint this timing delay precisely, which is unlikely achievable in this way, but rather to shift the traces by roughly the right amount to aid comparisons. The critical differences in timing across conditions are unaffected by this.

In order to use CPP amplitude modulations as additional constraints in the model fits to compare the explanatory power of neurally-informed models with and without drift rate bias (Figure 11B), we simply re-ran the behavioral fitting procedures but with a penalty term added to the G^2^ value to comprise the objective function. This penalty term was computed as the sum of squared differences between the percentage validity modulations (valid pre-response CPP amplitude with respect to the invalid condition) in the real and simulated CPP waveforms, summed across the three regimes, and weighted with a value logarithmically ranging between 0.01 and 10. Note that this particular method of ‘effectively’ constraining the CPP amplitude falling out of Monte Carlo simulations will necessarily result in worse fits to behavior considered alone because the fitting algorithm itself is increasingly de-emphasising the behavior in its objective function, and since it so strongly emphasises quantitative differences in ERP amplitudes it will be highly sensitive to any noise in those amplitudes. The purpose of this analysis was to examine how the relative performance of the model with versus without drift rate bias changed with emphasis on capturing CPP amplitude modulations.

## Acknowledgments

The authors thank Sarita Tamang and Genevieve Price for help with data collection. This work was supported by a research grant to S.P.K. and R.O.C. from the U.S. National Science Foundation under grant number BCS-1358955, a Career Development Award from Science Foundation Ireland to S.P.K. (15/CDA/3591), and a European Research Council (ERC) Starting Grant to RGOC under the European Union’s Horizon 2020 research and innovation programme (grant no. 638289). E.A.C. was supported by a Government of Ireland Postdoctoral Fellowship from the Irish Research Council (GOIPD/2017/1261).

## FIGURE SUPPLEMENTS

**Figure 2 - figure supplement 1:**
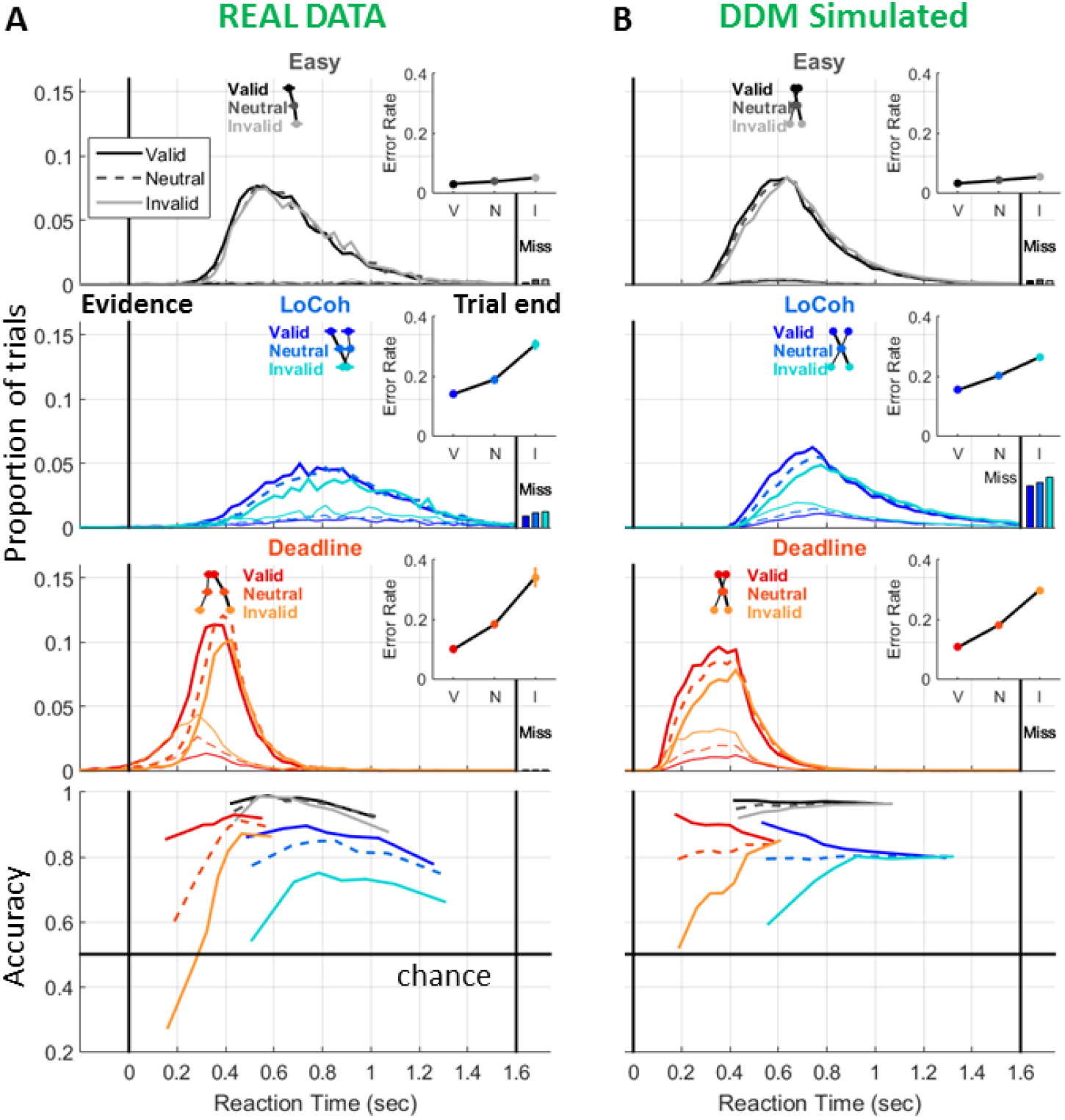
Same as figure 2 main figure but showing model with no drift rate biases allowed.

**Figure 5 - supplemental figure 1:**
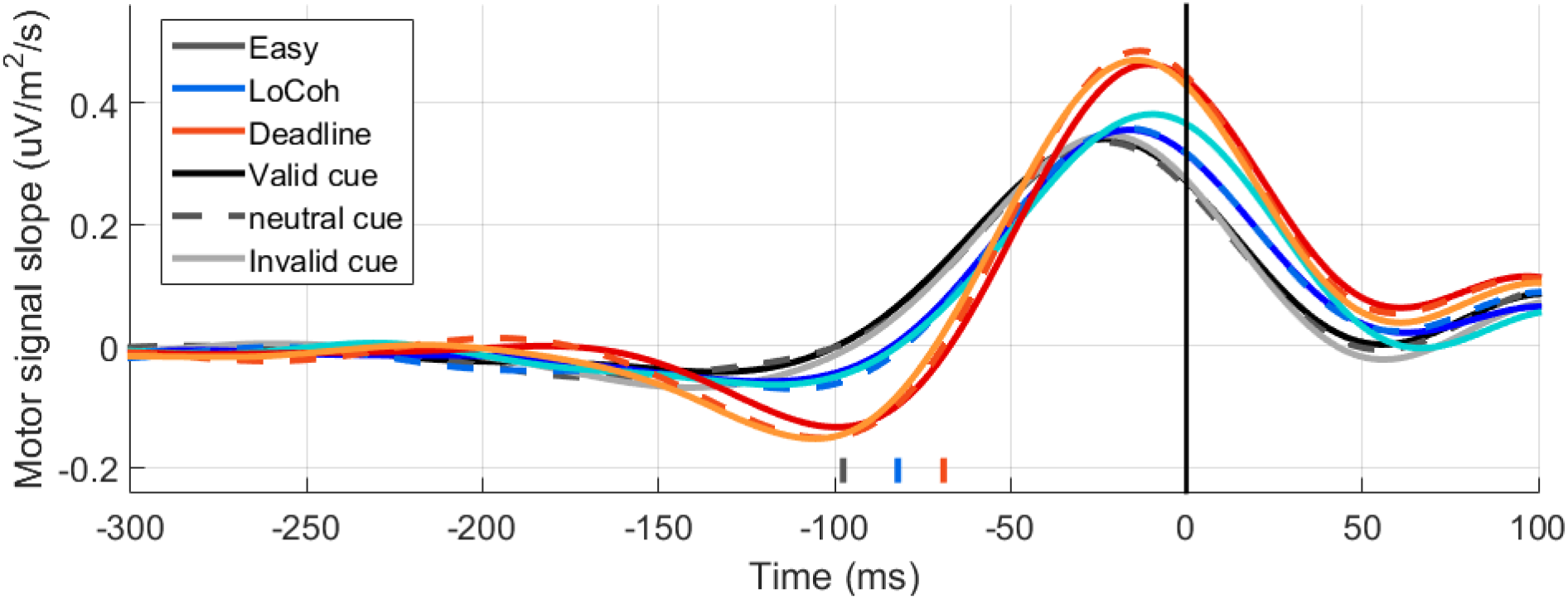
The first temporal derivative of the same motor signals as shown in Figure 5, quantifying momentary signal slope. The time point at which the first-derivative signal for each condition passes through zero, indicated below the waveforms, marks the inflection point for the motor signal, where negative-going motor preparation activity gives way to a sharp potential assumed to reflect motor execution.

**Figure 8 - figure supplement 1:**
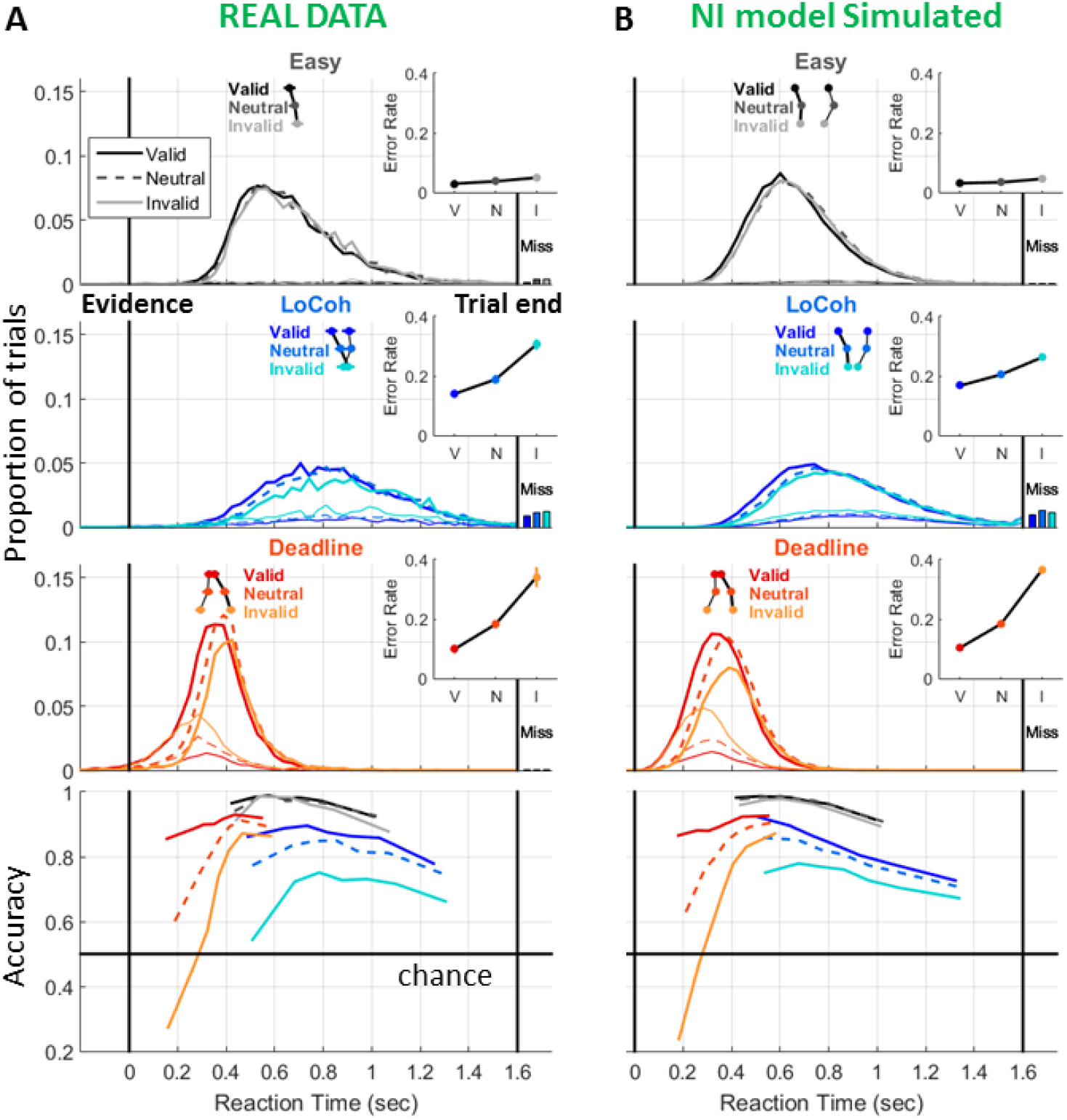
Same as figure 8 main figure but showing model with no drift rate biases allowed.

**Figure 9 - figure supplement 1:**
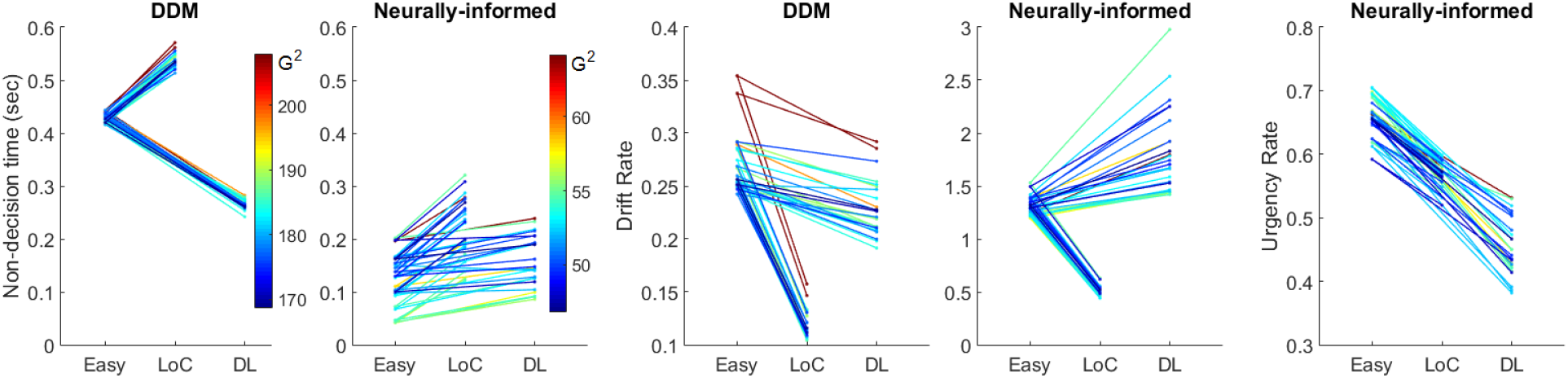
All 25 fits, even though each was started from an independent starting parameter vector originally and are of varying quality, agree on the qualitative trends across regimes for each parameter effect within each model. Bluer traces indicate lower G^2^ values (better fits), whereas redder indicate higher G^2^ (worse fits).

**Figure 10 - Supplemental figure 1:**
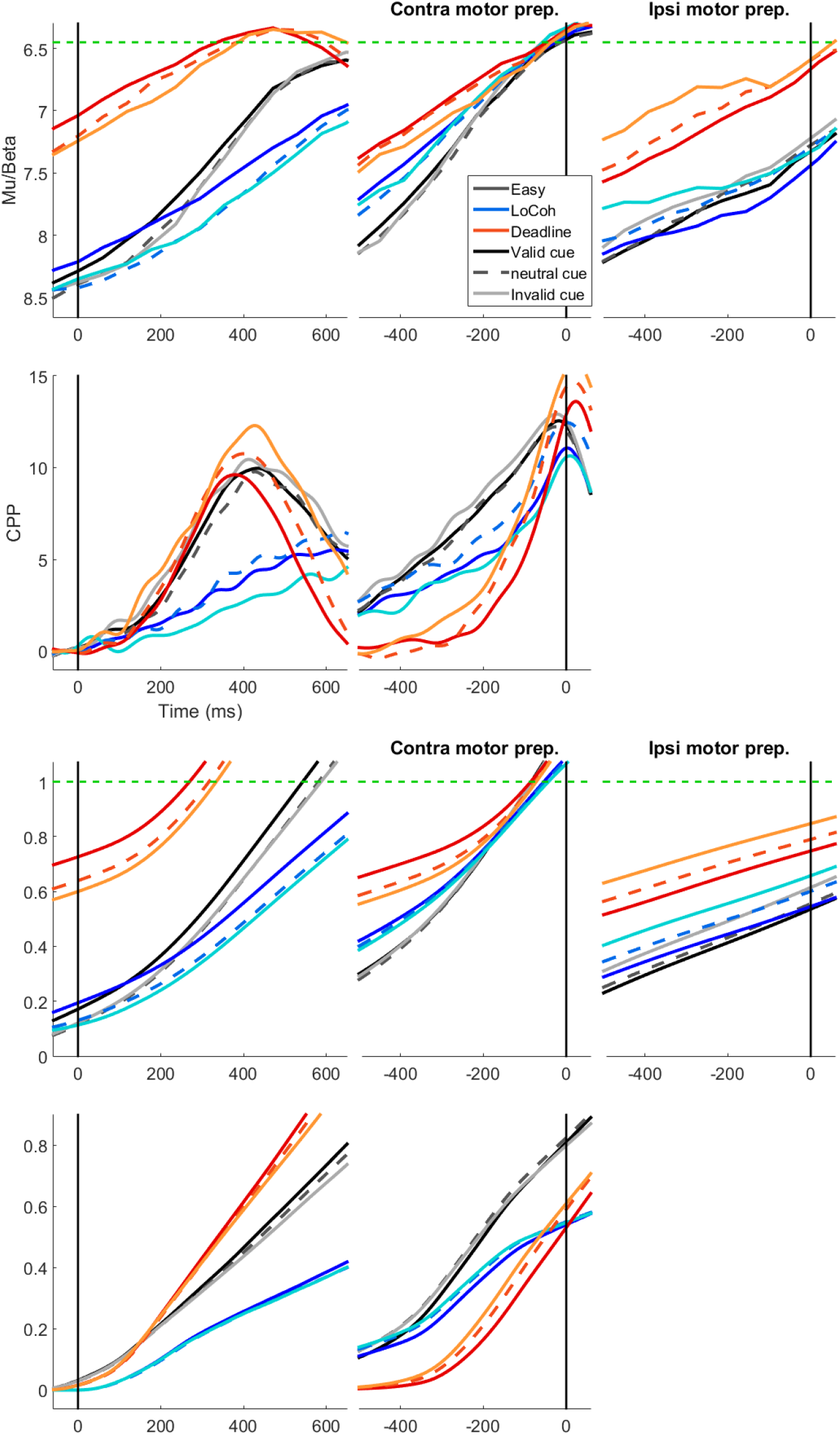
same real and simulated waveforms as main figure 10 but with each regime’s traces further broken out by prior cue validity. The simulations demonstrate another instance where an intuitive link between a model parameter and decision signal characteristic does not confer a reliable correspondence in trial-averaged data (Purcell and Palmeri 2017). Specifically, despite a clear drift rate bias in the model fit in the Deadline regime indicating that evidence is accumulated more steeply when validly than when invalidly cued, the simulated temporal slope of the accumulator signal ranks in the opposite way, due to other contingencies inherent in the model. This further highlights the importance of simulating decision variable dynamics rather than relying on correspondences between the model parameters themselves and the signal features. In this particular case, it demonstrates that neural evidence for the drift rate bias across regimes was better sought in CPP pre-response amplitude (Figure 11) rather than buildup rate.

